# Protection of double-Holliday junctions ensures crossing over during meiosis

**DOI:** 10.1101/2024.09.14.613089

**Authors:** Shangming Tang, Sara Hariri, Regina Bohn, John E McCarthy, Jennifer Koo, Mohammad Pourhosseinzadeh, Emerald Nguyen, Natalie Liu, Christopher Ma, Hanyu Lu, Monica Lee, Neil Hunter

## Abstract

Chromosomal linkages formed through crossover recombination are essential for accurate segregation of homologous chromosomes during meiosis^1^. The DNA events of recombination are linked to structural components of meiotic chromosomes^2^. Imperatively, the biased resolution of double-Holliday junction intermediates (dHJs) into crossovers^3,4^ occurs within the synaptonemal complex (SC), the meiosis-specific structure that mediates end-to-end synapsis of homologs during the pachytene stage^5,6^. However, the SC’s roles in crossover-specific dHJ resolution remains unclear. Here, we show that key SC components function through dependent and interdependent relationships to protect dHJs from aberrant “dissolution” into noncrossover products. Conditional ablation experiments reveal that cohesin, the core of SC lateral elements, is required to maintain both synapsis and dHJ-associated crossover recombination complexes (CRCs) during pachytene. The SC central region transverse-filament protein is also required to maintain CRCs. Reciprocally, stability of the SC central region requires the continuous presence of CRCs, thereby coupling synapsis and desynapsis to dHJ formation and resolution. However, dHJ protection and maintenance of CRCs can occur without end-to-end homolog synapsis mediated by the central element of the SC central region. We conclude that local ensembles of SC components are sufficient to enable crossover-specific dHJ resolution and thereby ensure the linkage and segregation of homologous chromosomes.

---

During meiotic prophase I, cohesin complexes connect sister-chromatids and mediate their organization into linear arrays of chromatin loops tethered to a common axis^2,5,7,8^. These cohesin-based axes define interfaces for the pairing and synapsis of homologous chromosomes that culminates in the formation of synaptonemal complexes (SCs), tripartite structures comprising the two juxtaposed homolog axes, now called lateral elements, connected by a central lattice of transverse filaments^5,6^. Extension of this lattice to achieve full synapsis requires an additional central-element complex (see **Extended Data Fig. 1a**)^5,6,9^. Meiotic recombination facilitates pairing and synapsis between homologous chromosomes, and then connects them via crossing over, which is necessary for accurate segregation during the first meiotic division^1^. To this end, the DNA events of recombination are physically and functionally linked to underlying chromosome structures^2^. The protein complexes that catalyze DNA double-strand breaks (DSBs) and subsequent strand exchange are tethered to homolog axes. Ensuing joint-molecule intermediates and their associated recombination complexes interact with the SC central region. A subset of recombination events is assigned a crossover fate with a tightly regulated distribution, ensuring that each chromosome pair receives at least one^2^. At designated sites, nascent joint molecules mature into double Holliday junctions (dHJs) and then undergo biased resolution specifically into crossovers^3,4^, all within the context of the SC central region and associated crossover recombination complexes (CRCs). Post-synapsis roles of SC components in crossing over remain unclear, particularly whether they function after dHJ formation to facilitate crossover-specific resolution.

## Cohesin is required for crossover-biased dHJ resolution

To address the role of SC lateral elements in crossover-specific dHJ resolution, the cohesin core was ablated using the auxin-inducible degron (AID) system^10^ to conditionally degrade Rec8 (the meiosis-specific Kleisin subunit) at the time of dHJ resolution (**Fig. 1a**). Real-time inactivation of Rec8-cohesin circumvents severe early meiotic defects of cohesin mutants in the formation and processing of DSBs (**Extended Data Fig. 2a–d**)^11,12^. In all experiments, cell cultures were synchronized at the resolution transition using an estradiol-inducible allele of *NDT80* (*NDT80-IN*) that reversibly arrests cells at the pachytene stage, in which chromosomes are fully synapsed and dHJs are poised for resolution^13^.

**Fig. 1.**
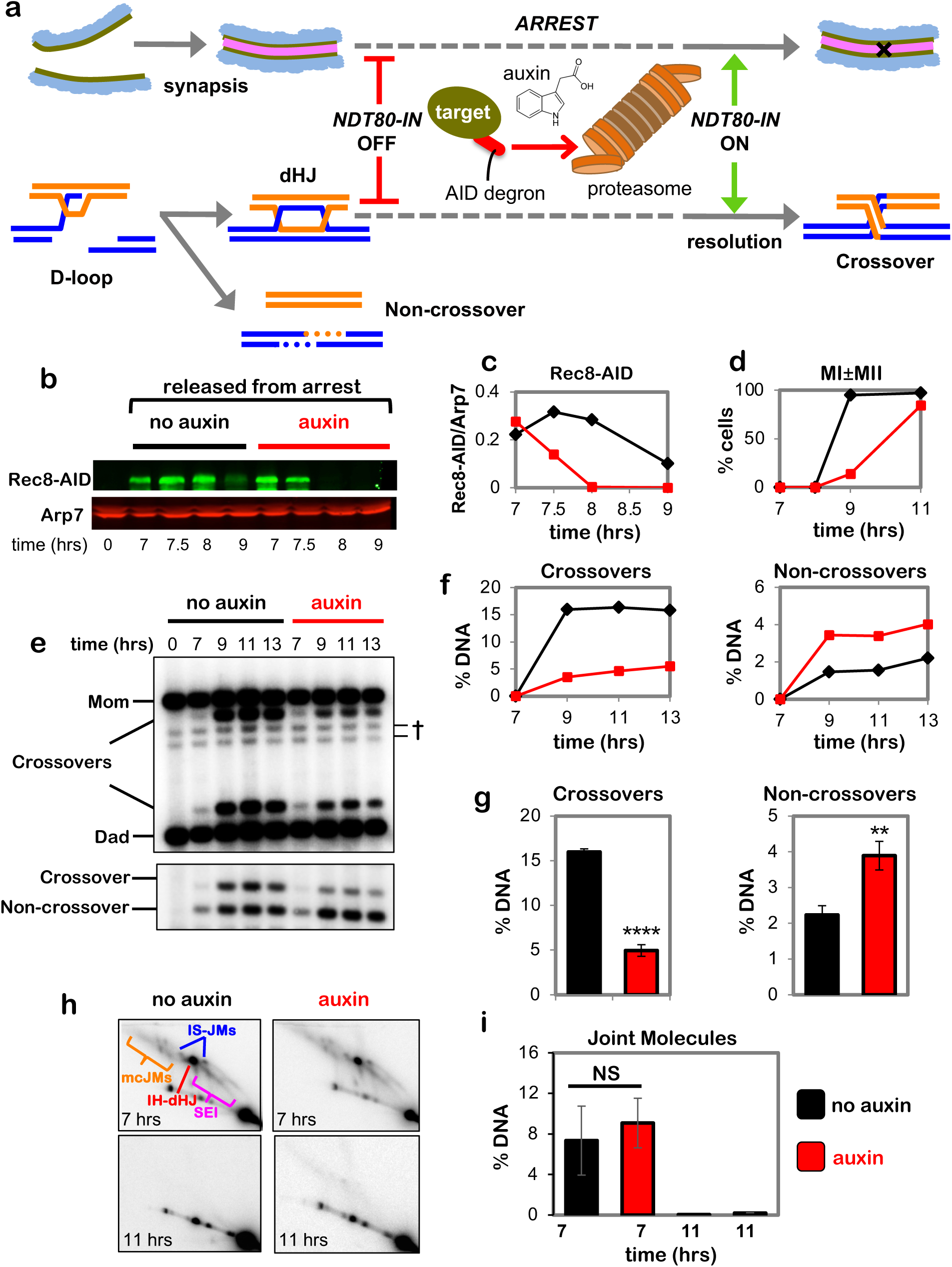
Rec8-cohesin is required for crossover-specific dHJ resolution. **a,** Experimental strategy. Top row: homolog synapsis with chromatin in blue, green homolog axes, and pink SC central region. The **X** in the final cartoon indicates a crossover formed after Ndt80 is expressed. Middle row: cell synchronization at the dHJ resolution transition using the inducible *NDT80-IN* allele; and conditional degradation of target proteins using the AID system. Bottom row: DNA events of meiotic recombination. Only the two chromosomes engaged in recombination are shown. **b,** Western analysis of Rec8-AID from subcultures with or without the addition of auxin at 7 hours. Arp7 is used as a loading control. **c,** Quantification of Rec8-AID level from the experiment shown in panel **b**. **d,** Quantification of nuclear divisions (MI±MII, cells with two and four nuclei) from *REC8-AID* subcultures with or without auxin. **e,** Representative 1D-gel Southern analysis of crossover (upper panel) and non-crossover (lower panel) formation at the *HIS4::LEU2* recombination hotspot from *REC8-AID* subcultures with or without auxin. ^†^ cross-hybridizing background bands. **f,** Quantification of crossovers and non-crossovers levels from the experiments shown in panel **e**. **g,** Quantification of final levels of crossovers and non-crossovers at 11 hrs from *REC8-AID* subcultures with or without auxin (mean ± SD, 3 independent experiments, \*\*\*\**P*<0.0001, \*\**P*=0.0038. two-tailed unpaired *t*-test). **h,** Representative 2D gel Southern analysis of joint molecules from *REC8-AID* subcultures with or without auxin, before and after release from arrest at 7 and 11 hrs, respectively. The upper left panel highlights joint-molecule species: SEI, single end invasion; IH-dHJ, inter-homolog double Holliday junction; IS-JMs, inter-sister joint molecules; mc-JMs, 3- and 4-chromatid joint molecules. **i,** Quantification of total joint molecule levels from experiments represented in panel **h** (mean ± SD, *n=*3 independent experiments, NS, not significant, *P*=0.4561, two-tailed unpaired *t*-test).

Auxin and estradiol were added simultaneously to degrade Rec8-AID while releasing cells from pachytene arrest and triggering dHJ resolution (**Fig. 1b–d**). Without auxin, Rec8-AID levels remained high until cells had completed meiotic divisions (MI±MII); but with auxin, Rec8-AID was completely degraded within 60 mins and meiotic divisions were delayed by ∼2 hrs (**Fig. 1c** and **1d**). Crossing over at the *HIS4::LEU2* DSB hotspot^14^ was reduced by 70% following Rec8-AID degradation, accompanied by a 43% increase in noncrossover products (**Fig. 1e–g**; note that our assay reports a subset of noncrossover gene-conversion products, not absolute levels of noncrossovers^14^). This reciprocal change in levels of crossover and noncrossover products, and the comparable kinetics of their formation with and without auxin (**Fig. 1e,f**), suggest that dHJs are efficiently resolved when Rec8-AID is degraded but their resolution fate is reversed. This inference was confirmed by 2D-gel electrophoresis and Southern blotting (**Fig. 1h** and **1i**). Thus, Rec8-based cohesin is required after dHJ formation to facilitate crossover-specific resolution. Degradation of core cohesin subunit, Smc3, confirmed that the cohesin complex, not just Rec8, is required (**Extended Data Fig. 2e–h**). However, the mitotic Kleisin, Mcd1^RAD21^, is not required for crossover-specific dHJ resolution indicating that this function is specific to Rec8-cohesin (**Extended Data Fig. 2i,j**). We also showed that separase, Esp1, does not influence crossover-specific resolution (**Extended Data Fig. 2i**), consistent with Yoon et al. ^15^ who showed that a separase-resistant allele of Rec8 does not affect crossing over.

## Cohesin and Smc5/6 function in distinct resolution pathways

Two classes of meiotic crossovers are distinguished by their dependencies on joint-molecule resolving enzymes: class-I crossovers depend on the crossover-specific dHJ resolvase defined by the endonuclease MutLγ (Mlh1-Mlh3)^3,4,16–18^; while minority class-II crossovers require structure-selective endonucleases, primarily Mus81-Mms4^Eme1^ ^3,19–21^. Epistasis analysis revealed that cohesin and MutLγ act in the same crossover resolution pathway (**Fig. 2a,b**); while Mus81-Mms4^Eme1^ acts in a parallel pathway (**Fig. 2c,d**, in these experiments Mms4-AID was degraded at the resolution transition in a *yen1Δ* background in which backup resolvase Yen1 was deleted). Crossover levels were indistinguishable following Rec8-AID degradation in the presence or absence of MutLγ (*mlh3Δ* mutation; **Fig. 2b**). Noncrossover formation in *REC8-AID mlh3Δ* strains was reduced to control levels suggesting that MutLγ influences whether products formed in the absence of cohesin are associated with gene conversion of the SNP employed to detect noncrossovers^14^.

**Fig. 2.**
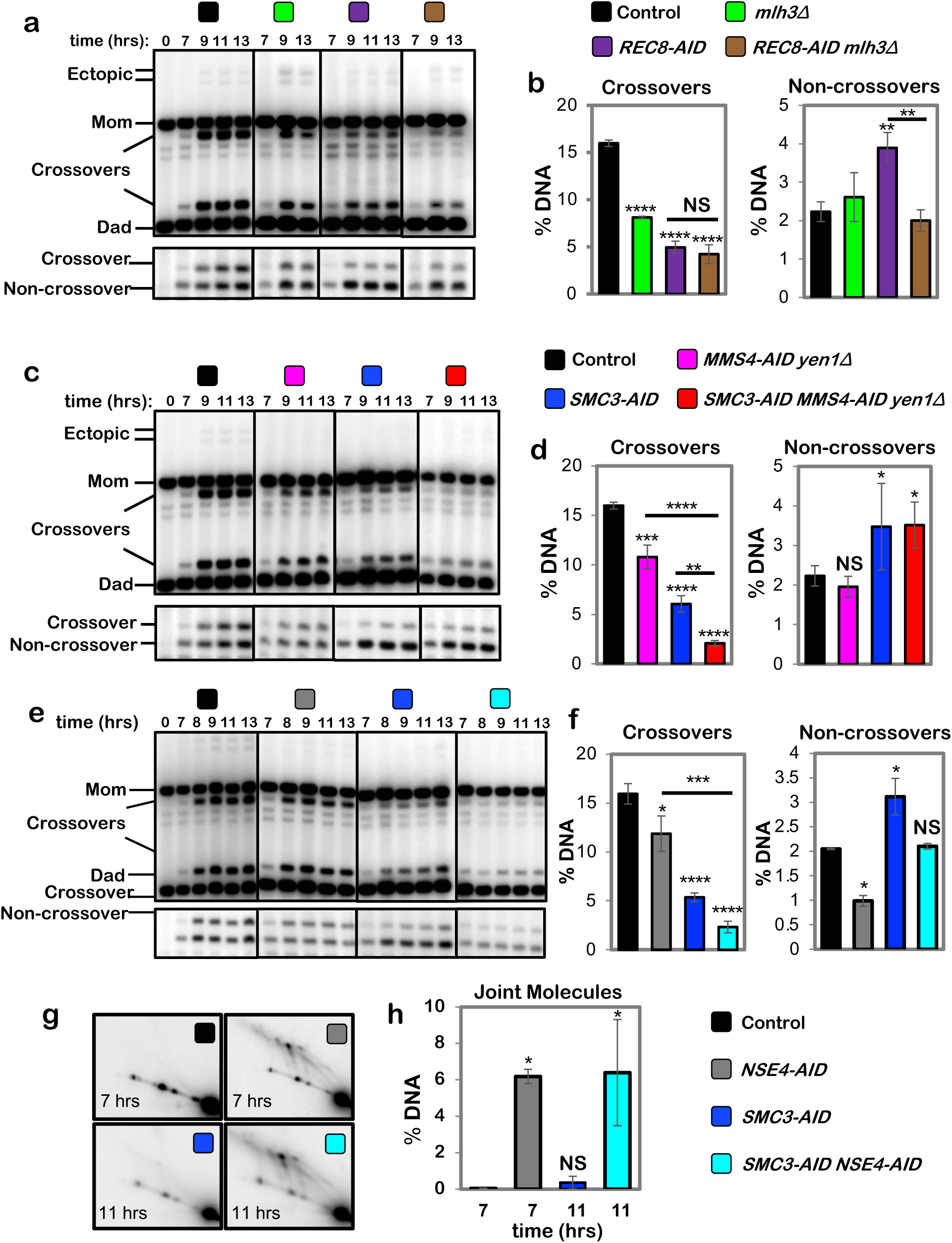
Rec8-cohesin and Smc5/6 define distinct pathways of joint-molecule resolution. In all experiments, cells were released from *NDT80-IN* arrest. **a,** Representative 1D-gel Southern analysis of crossover (upper panel) and non-crossover (lower panel) formation in control (*REC8-AID* no auxin), *mlh3*Δ, *REC8-AID* (with auxin) and *REC8-AID mlh3*Δ (with auxin) strains. **b,** Final levels of crossovers and non-crossovers at 13 hrs from the indicated strains (mean ± SD, 3 independent experiments). Statistical comparisons with control unless indicated. Dunnett’s multiple comparisons test, \*\*\*\**P*<0.0001, NS, not significant *P*=0.5281, \*\**P*=0.0034 (*REC8-AID* vs. control), \*\**P*=0.0026 (*REC8-AID* vs. *REC8-AID mlh3*Δ). **c,** Representative 1D-gel Southern analysis of crossover (upper panel) and non-crossover (lower panel) formation in control, *MMS4-AID yen1*Δ (with auxin), *SMC3-AID* (with auxin) and *SMC3-AID MMS4-AID yen1*Δ (with auxin) strains. **d,** Final levels of crossovers and non-crossovers at 11 hrs from the indicated strains (mean ± SD, 3 independent experiments. Statistical comparisons with control unless indicated. For crossovers, Tukey’s multiple comparisons test was performed; for non-crossovers, Dunnett’s multiple comparisons test was performed. \*\*\*\**P*<0.0001, \*\*\**P*=0.0003, \*\**P*=0.0066, \**P*= 0.0385 (control vs. *SMC3-AID*), \**P*=0.0488 (control vs. *SMC3-AID MMS4-AID yen1*Δ.), NS, not significant *P*=0.7806). **e,** Representative 1D-gel Southern analysis of crossover (upper panel) and non-crossover (lower panel) formation in control, *NSE4-AID* (with auxin), *REC8-AID* (with auxin) and *REC8-AID NSE4-AID* (with auxin) strains. **f,** Final levels of crossovers and non-crossovers at 11 hrs from the indicated strains (mean ± SD, 3 independent experiments). Statistical comparisons with control unless indicated. For crossovers, Tukey’s multiple comparisons test was performed; for non-crossovers, Dunnett’s multiple comparisons test was performed. \*\*\*\**P*<0.0001, \*\*\**P*=0.0003, \**P*= 0.0262 (control vs, *NSE4-AID* for crossover comparison), \**P*= 0.0464 (control vs. SMC3-AID for non-crossover comparison), NS, not significant, *P*=0.1148 (control vs. *NSE4-AID*), *P* >0.9999 (control vs. *SMC3-AID NSE4-AID*). **g,** Representative 2D gel Southern analysis of joint molecules from control, *NSE4-AID* (with auxin), *SMC3-AID* (with auxin) and *SMC3-AID NSE4-AID* (with auxin) strains. **h,** Quantification of total joint-molecule levels from the indicated strains. (mean ± SD, 3 independent experiments). Dunnett’s multiple comparisons test with control: \**P*=0.0164 (control vs. *NSE4-AID*), \**P*=0.0097 (control vs. *SMC3-AID NSE4-AID*, NS, not significant, *P*=0.9954).

Mus81-Mms4^Eme1^ works in conjunction with a second SMC complex, Smc5/6 ^20–23^. Consistently, degradation of Nse4-AID (an essential subunit of Smc5/6) at the resolution transition reduced crossing over to the same extent as when Mus81-Mms4^Eme1^ was inactivated by degrading Mms4-AID (**Fig. 2c–f**). However, while noncrossovers were not reduced when Mus81-Mms4^Eme1^ was inactivated, Nse4-AID degradation also reduced noncrossovers 2.1-fold (**Fig. 2e** and **2f**), indicating that Smc5/6 controls an additional, Mus81-Mms4^Eme1^-independent pathway of noncrossover formation most likely by regulating a subpopulation of the Sgs1-Top3-Rmi1 complex (**Extended Data Figs. 1 and 3**).

Confirming that Smc5/6 and cohesin act in independent pathways of resolution, simultaneous degradation of Nse4-AID and Smc3-AID resulted in an additive reduction in crossing over; and noncrossover levels that are the combination of those observed when Nse4-AID and Smc3-AID were degraded individually (**Fig. 2e** and **2f**; co-degradation of Nse4-AID and Rec8-AID gave analogous results, **Extended Data Fig. 4**). Further distinguishing these two pathways, a subset of joint molecules remained unresolved when Nse4-AID was degraded alone, while resolution remained efficient when cohesin was inactivated (via degradation of either Rec8-AID or Smc3-AID; **Figs. 1g,h** and **2g,h** and **Extended Data Fig. 4**); that is, Smc5/6 is essential for the resolution of a subset of joint molecules into both crossovers and noncrossovers, while Rec8-cohesin specifically promotes crossover-specific dHJ resolution, but is not required for resolution *per se* (see **Extended Data Fig. 4c–f**).

## Rec8-cohesin is required to maintain synapsis and crossover recombination complexes during pachytene

To begin to understand how cohesin facilitates crossover-specific dHJ resolution, Rec8-AID was degraded while pachytene arrest was maintained (no induction of *NDT80-IN*) and chromosomes were analyzed by immunostaining for markers of homolog axes (Rec8 and Red1), SC central region (Zip1), and crossover recombination complexes (CRCs; Msh5 and Zip3)^24^(**Figure 3a**). In no-auxin controls, linear Rec8 and discontinuous Red1 staining structures colocalized; and synapsed homologs (indicated by lines of Zip1 staining) were decorated with foci of Msh5 and Zip3 (**Fig. 3a**). One hour after auxin addition, Rec8 structures were lost and characteristic features of pachytene chromosomes disappeared: SCs disassembled and Zip1 staining was now confined to a few foci and larger structures resembling polycomplexes that are diagnostic of defective synapsis; numbers of Red1 structures were reduced 2-fold, consistent with cohesins’ core function in organizing homolog axes^25^ and directly recruiting Red1^26^; and CRCs dissociated indicated by loss of Msh5 and Zip3 foci (**Fig. 3a** and **3b**). Thus, cohesin is required to maintain the integrity of pachytene chromosome structures, consistent with analysis of meiotic cohesin in *C. elegans*^27^.

**Fig. 3.**
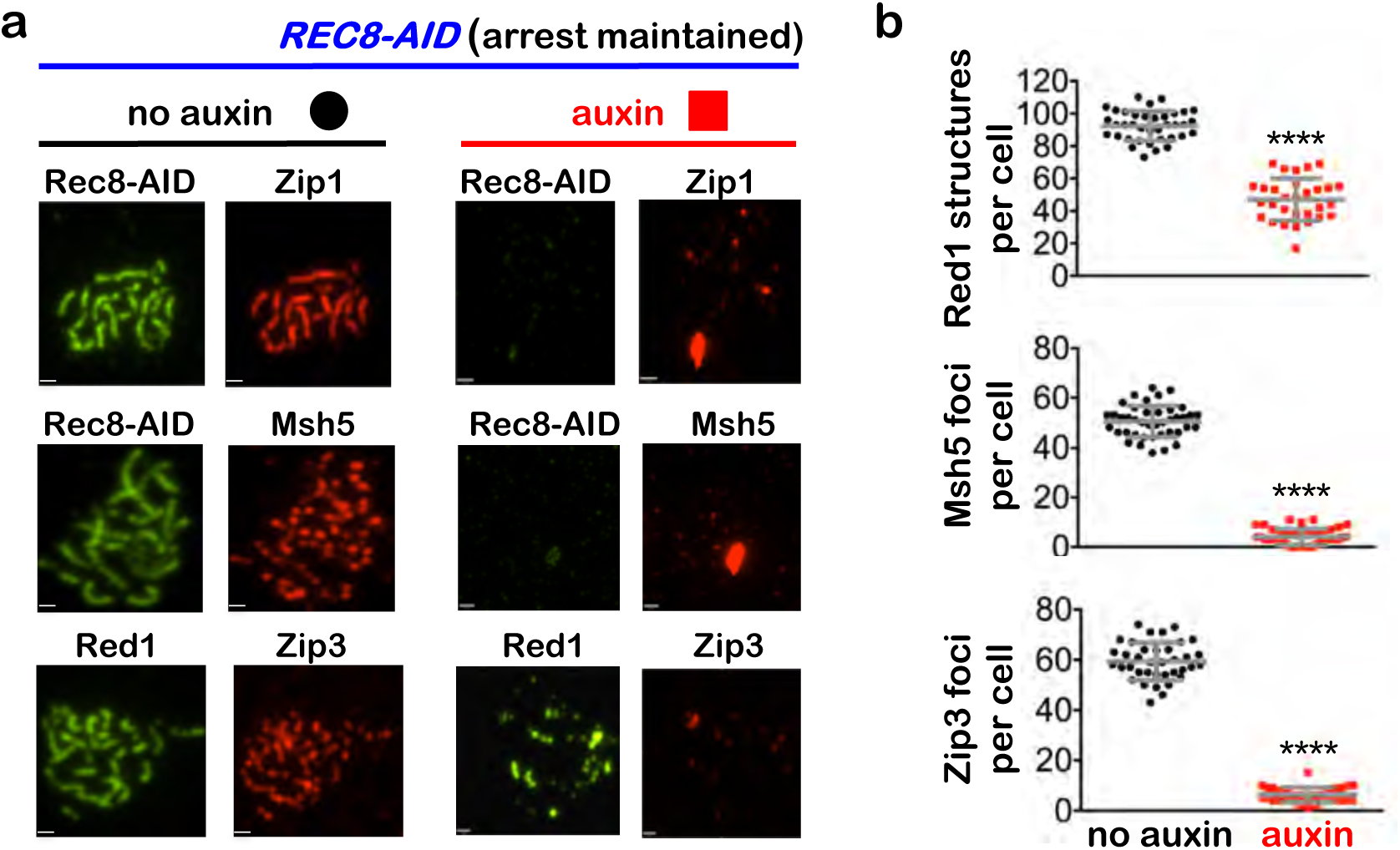
Rec8-cohesin is required to maintain synaptonemal complexes and crossover-specific recombination complexes. **a,** Representative images of surface-spread meiotic nuclei from *REC8-AID* cultures in which pachytene-arrest was maintained. Cells were sampled 1 hr after the addition of auxin or DMSO vehicle at 7 hrs, and immunostained for the indicated markers. Scale bars = 1 μm. **b,** Quantification of Red1, Msh5 and Zip3 immunostaining structures from experiments represented in **a**. 40-60 nuclei were counted in each case. Error bars represent SD. Unpaired two-tailed *t* test. \*\*\*\**P*<0.0001.

## Interdependent functions of the SC components are required for crossover-specific dHJ resolution

To discern which cohesin-dependent feature(s) of pachytene chromosomes is important for crossover-specific dHJ resolution, SC transverse-filament protein Zip1^28,29^ and pro-crossover factor MutSγ were inactivated as cells were released from pachytene arrest (**Fig. 4a,b**). MutSγ (a complex of Msh4 and Msh5) binds and stabilizes nascent joint molecules to promote homolog synapsis and dHJ formation^30–32^. Both Zip1-AID and Msh4-AID degradation caused phenotypes similar to those resulting from loss of cohesin, i.e. reduced crossovers and increased noncrossovers (**Fig. 4c,d**), indicating continued roles after dHJs have formed to maintain their crossover resolution fate.

**Figure 4.**
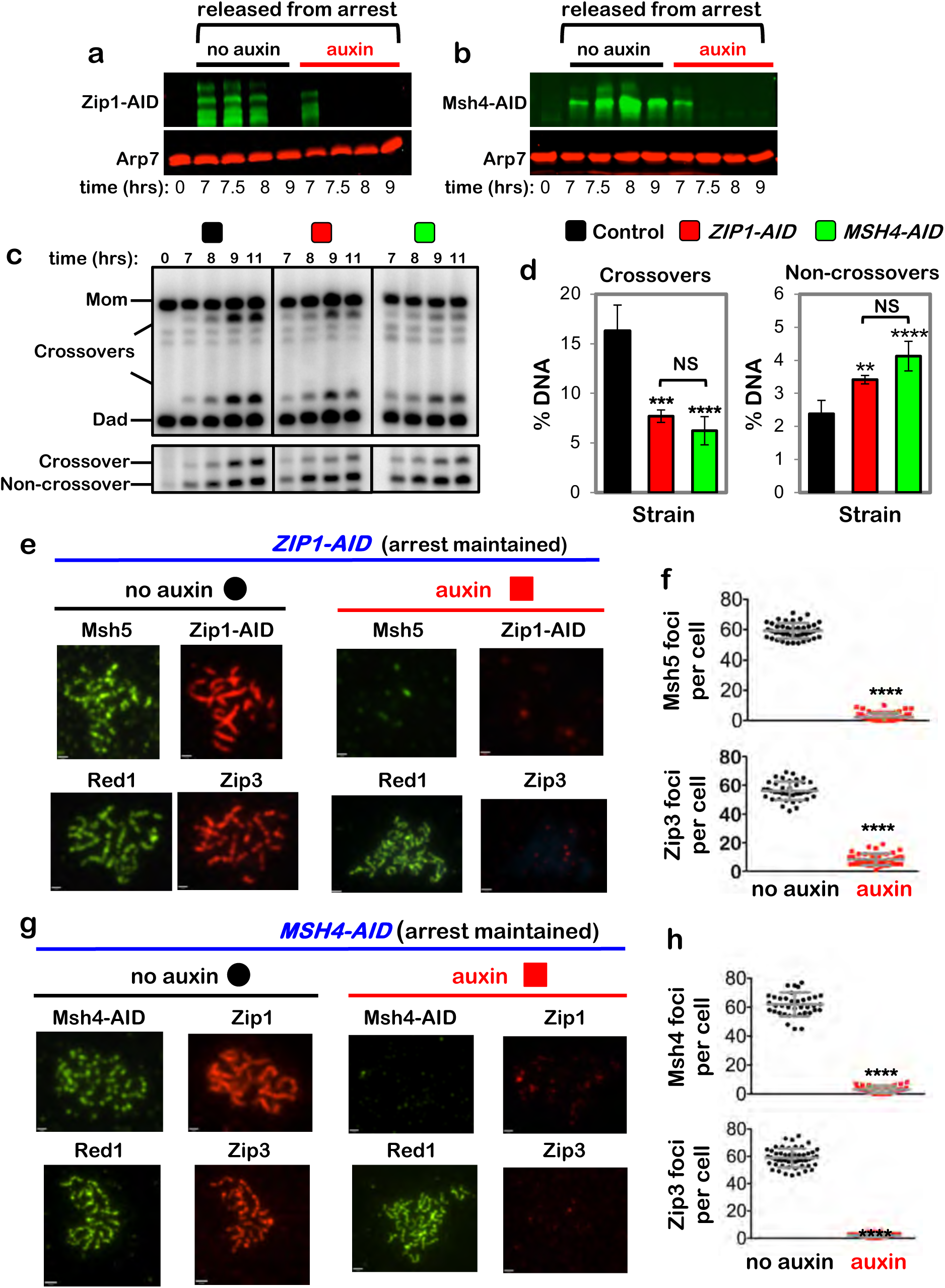
Zip1 and MutSγ are required for crossover-specific dHJ resolution. **a,** Western blot analysis of Zip1-AID from subcultures with or without the addition of auxin at 7 hours. Arp7 is a loading control. **b,** Western blot analysis of Msh4-AID from subcultures with or without the addition of auxin at 7 hours. Arp7 is used as a loading control. **c,** Representative 1D-gel Southern analysis of crossover (upper panel) and non-crossover (lower panel) formation in control, *ZIP1-AID* (with auxin) and *MSH4-AID* (with auxin) strains. **d,** Final levels of crossovers and non-crossovers at 11 hrs from the indicated strains. Error bars represent SD for three independent experiments. Statistical comparisons with control unless indicated. Tukey’s multiple comparisons test, \*\*\*\**P*<0.0001, \*\**P*=0.0099, NS, not significant, *P*=0.3353 (*ZIP1-AID* vs. *MSH4-AID* for crossover analysis), *P*=0.0763 (*ZIP1-AID* vs. *MSH4-AID* for non-crossover analysis). **e,** Representative images of surface-spread meiotic nuclei from pachytene-arrested *ZIP1-AID* cells sampled 1 hr after the addition of auxin or DMSO vehicle at 7 hrs, and immunostained for the indicated markers. Scale bars = 1 μm. **f,** Quantification of Msh5 and Zip3 immunostaining foci from the experiments represented in panel **e**. 40-60 nuclei were counted in each case. Error bars represent SD. Unpaired two-tailed *t* test, *P*<0.0001. **g,** Representative images of surface-spread meiotic nuclei from pachytene-arrested *MSH4-AID* cells sampled 1 hr after the addition of auxin or DMSO vehicle at 7 hrs, and immunostained for the indicated markers. **h,** Quantification of Msh4 and Zip3 immunostaining foci from the experiments represented in panel **g**. 40-60 nuclei were counted in each case. Error bars represent SD. 40-60 nuclei were counting for each set of data. In **e** and **g**, scale bars = 1 μm. Unpaired two-tailed *t* test, \*\*\*\**P*<0.0001.

This analysis suggests that cohesin may facilitate crossover-specific dHJ resolution by stabilizing the SC central region, which in turn stabilizes CRCs. This interpretation was tested by immunostaining the chromosomes of pachytene-arrested cells following degradation of Zip1-AID (**Fig.4e,f**) or Msh4-AID (**Fig.4g,h**), respectively. When Zip1-AID was degraded, homologs desynapsed but axis integrity was maintained as shown by robust Red1 localization (**Fig. 4e**). As predicted, CRCs marked by Msh5 and Zip3 foci were diminished (**Fig. 4e,f**). As Zip3 and Msh4 protein levels remained high in these cells (**Extended Data Fig. 5**), we infer that CRCs are disassembled when Zip1-AID is degraded. Unexpectedly, desynapsis occurred when Msh4-AID was degraded revealing that the maintenance of SCs during pachytene requires the continued presence of CRCs (**Fig. 4g**). Zip3 foci were also lost when Msh4-AID was degraded highlighting the interdependence between Holliday-junction binding (MutSγ) and regulatory (Zip3) components of CRCs (**Fig. 4g,h**).

## Pachytene chromosome structures protect dHJs from noncrossover resolution by the Sgs1^BLM^-Top3-Rmi1 complex

dHJs normally remain stable in pachytene-arrested *NDT80-IN* cells because expression of polo-like kinase Plk1^Cdc5^, which activates dHJ resolution, requires the Ndt80 transcription factor ^13,33^. However, dHJ levels decreased ∼3-fold when Rec8-AID was degraded while maintaining pachytene arrest (**Fig. 5b**,**c**) implying that pachytene chromosome structures protect dHJs from aberrant resolution via a Plk1^Cdc5^-independent resolution activity. A good candidate for this activity is Sgs1^BLM^-Top3-Rmi1 (STR), the budding yeast ortholog of human BLM complex BLM-TOPIIIα-RMI1/2, and a robust decatenase enzyme that can “dissolve” dHJs specifically into noncrossover products^34–38^. Confirming this prediction, dHJs were stabilized when Rec8-AID and Top3-AID were simultaneously degraded (**Fig. 5a–c**; co-degrading Smc3-AID and Top3-AID gave very similar results, **Extended Data Fig. 6**). Moreover, dHJ resolution following Rec8-AID degradation alone produced noncrossover products that did not form when Top3-AID was also degraded (**Fig. 5d,e**).

**Fig, 5.**
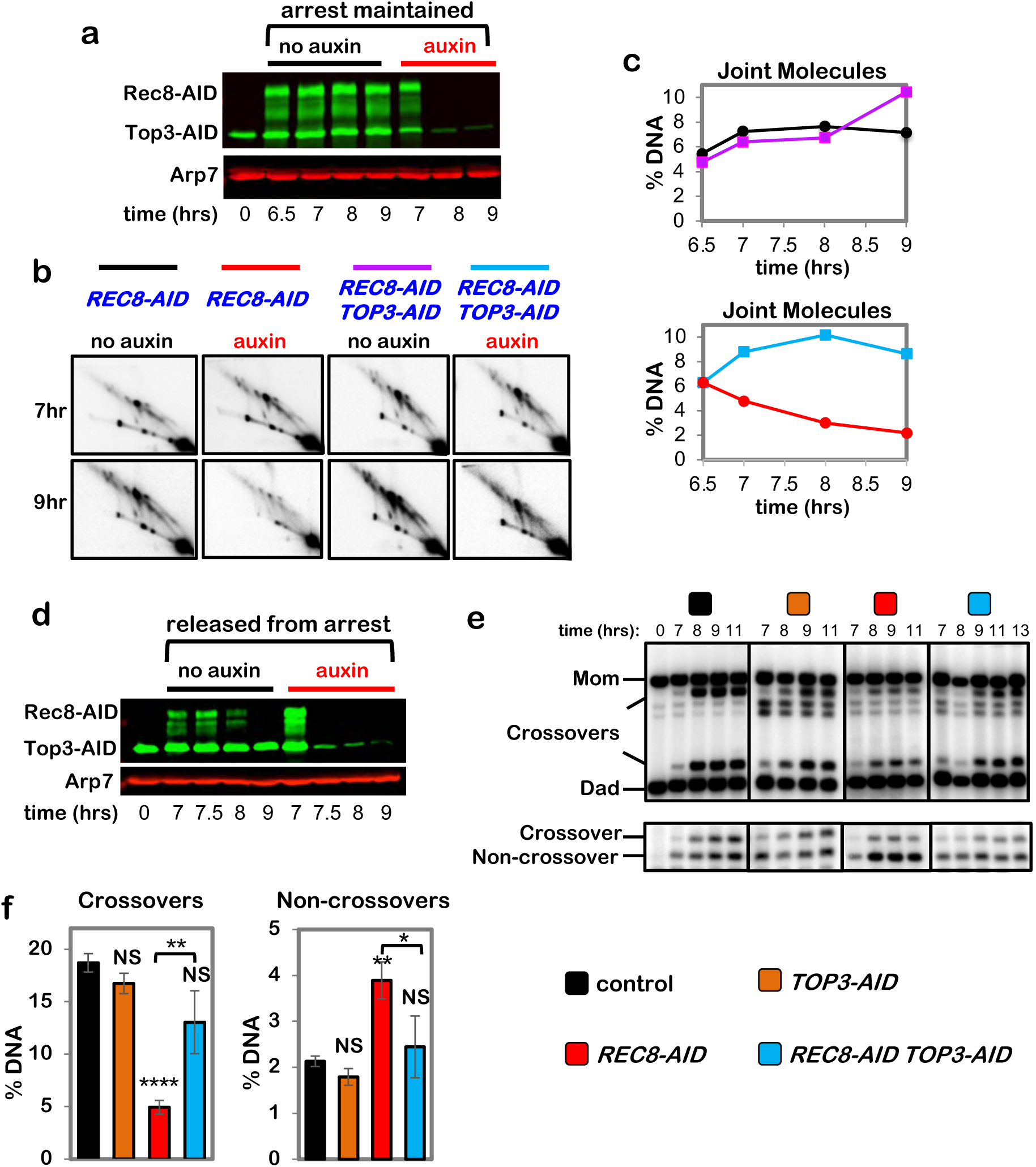
Rec8-cohesin protects double-Holliday junctions from aberrant resolution mediated by the BLM/STR complex. **a,** Western blot analysis of Top3-AID and Rec8-AID from subcultures with or without the addition of auxin at 7 hours while maintaining pachytene arrest. Arp7 is a loading control. The degradation-resistant fraction of Top3-AID is mitochondrial. **b,** Representative 2D gel Southern analysis of joint molecules from *REC8-AID* and *REC8-AID TOP3-AID* strains with and without the addition of auxin. **c,** Quantification of total joint molecule levels from the indicated strains. **d,** Western blot analysis of Top3-AID and Rec8-AID from subcultures with or without the addition of auxin at 7 hours with release from pachytene arrest. **e,** Representative 1D-gel Southern analysis of crossover (upper panel) and non-crossover (lower panel) formation in control, *TOP3-AID* (with auxin), *REC8-AID* (with auxin) and *REC8-AID TOP3-AID* (with auxin) strains. **f,** Final levels of crossovers and non-crossovers at 11 hrs from experiments represented in **g**. Error bars represent SD for three independent experiments. Statistical comparisons with control unless indicated. Tukey’s multiple comparisons test, \*\*\*\**P*<0.0001, \*\**P*=0.0011 (*REC8-AID* vs. *REC8-AID TOP3-AID* for crossover comparison), NS, not significant, *P*=0.8881 (control vs. *TOP3-AID* for crossover comparison), NS, not significant, *P*=0.0.1516 (control vs. *REC8-AID TOP3-AID* for crossover comparison), \*\**P*=0.0041 (control vs. REC8-AID for non-crossover comparison), \**P*=0.0156, NS, not significant, *P*=0.5131 (control vs. *TOP3-AID* for non-crossover comparison), *P*= 0.9200 (control vs. *REC8-AID TOP3-AID* for non-crossover comparison).

The stability of dHJs in pachytene-arrested cells following co-degradation of Rec8-AID and Top3-AID indicates that resolution is once again dependent on Plk1^Cdc5^ and suggests that the crossover defect resulting from Rec8-AID degradation may be rescued. Indeed, expression of *NDT80-IN* while simultaneously degrading Rec8-AID and Top3-AID resulted in a 2.6-fold increase in crossovers relative to degradation of Rec8-AID alone, while noncrossovers decreased 1.6-fold (**Fig. 5f–h**; co-degradation of Rec8-AID and Sgs1-AID gave similar results, **Extended Data Fig. 7**). Moreover, stabilization of dHJs and partial rescue of the crossover defects resulting from Zip1-AID or Msh4-AID degradation were also seen when Top3-AID was simultaneously degraded (**Extended Data Figs. 8 and 9**). Partial restoration of crossover levels is likely explained by our observation that when STR function is ablated, essentially all resolution is mediated by the Smc5/6–Mus81-Mms4 (**Extended Data Fig. 3**).

We wondered whether stabilizing dHJs also rescues the cytological defects resulting from degradation of Rec8-AID, Zip1-AID, or Msh4-AID. To this end, Top3-AID was co-degraded with Rec8-AID, Zip1-AID or Msh4-AID in pachytene-arrested cells, and chromosomes were immunostained for markers of synapsis (Zip1), and CRCs (Msh4 and Zip3)(**Extended Data Fig. 10**). In each case, cytological phenotypes were largely indistinguishable from those observed when Rec8-AID, Zip1-AID, or Msh4-AID were degraded alone. SCs and CRCs disassembled following co-degradation of Rec8-AID and Top3-AID (**Extended Data Fig. 10a,b**); CRCs dissociated when Zip1-AID and Top3-AID were co-degraded (**Extended Data Fig. 10c,d**); and SCs disassembled following co-degradation of Msh4-AID and Top3-AID (**Extended Data Fig. 10e,f**). Thus, maintaining inter-homolog DNA connections by stabilizing dHJs does not bypass the interdependencies between cohesin, SCs, and CRCs.

We conclude that the key components of pachytene chromosomes – cohesin-based homolog axes, SC transverse filaments, and CRCs – protect crossover-designated dHJs from being aberrantly dissolved into noncrossovers by the STR complex (see **Extended Data Fig. 1b**).

## Full synapsis is not essential for crossover-specific dHJ resolution

High levels of crossing over can occur without end-to-end homolog synapsis in cells that lack the SC central element, which is required to mature and extend the transverse filament lattice to achieve full synapsis^5,6,9,39^. To determine whether maintenance of full synapsis is required for crossover-specific dHJ resolution, central-element protein Ecm11^9^ was ablated as cells were released from pachytene arrest (**Fig. 6a,b**). Crossover and noncrossover levels were indistinguishable from no-auxin controls when Ecm11-AID was degraded, sharply contrasting phenotypes seen following loss of Rec8, Zip1, or Msh4 (**Figs. 1 and 4**). Moreover, in arrested *NDT80-IN* cells, dHJs remained stable following Ecm11-AID degradation (**Fig. 6c,d**).

**Fig. 6.**
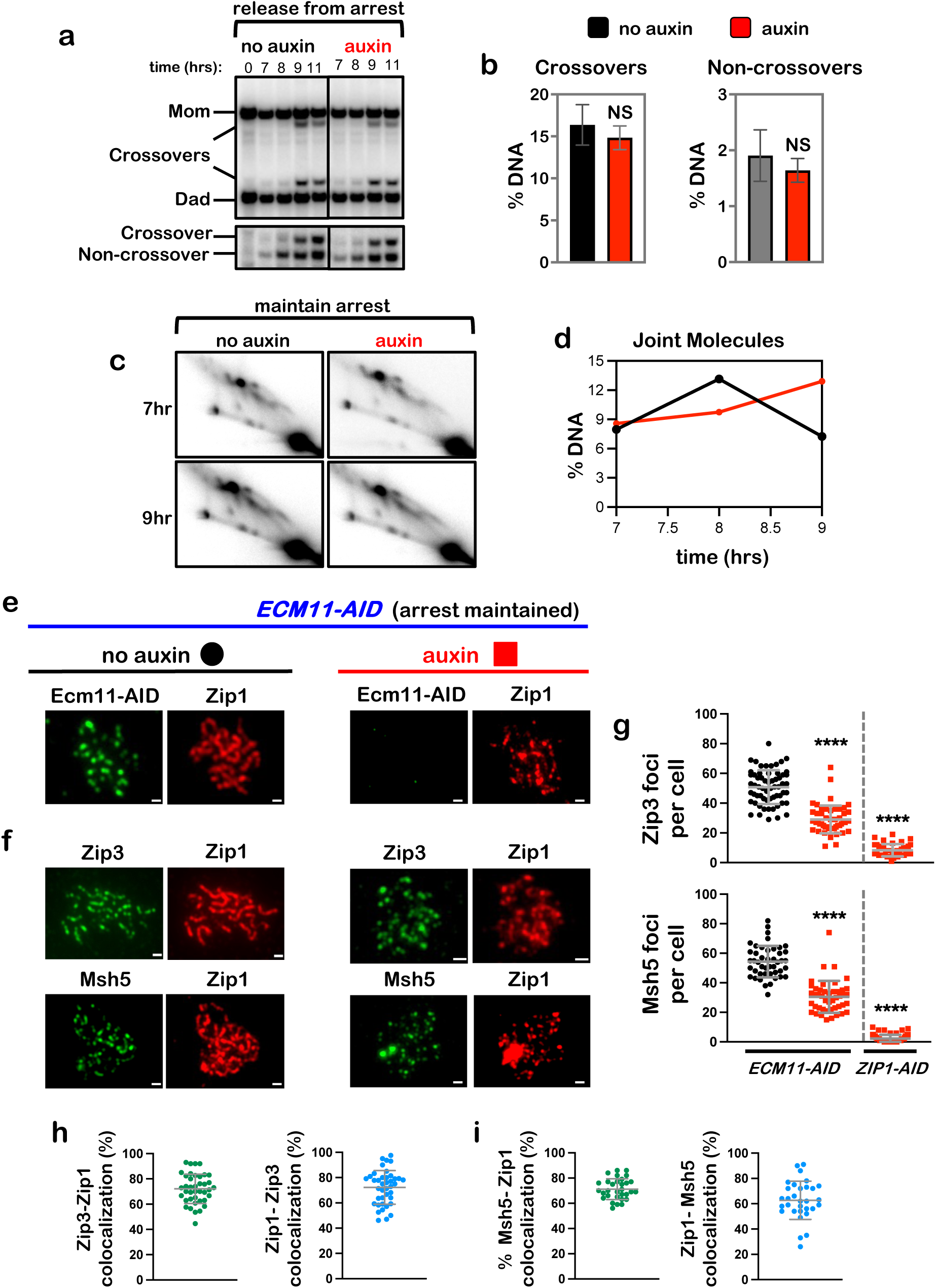
Full synapsis is not essential for crossover-specific dHJ resolution. **a,** Representative 1D-gel Southern analysis of crossover (upper panel) and non-crossover (lower panel) formation from *ECM11-AID* subcultures with or without auxin. NS, not significant, unpaired two-tailed *t*-test. **b,** Final levels of crossovers and non-crossovers at 11 hrs from the experiments represented in panel **a**. Error bars represent SD for three independent experiments. **c**, Representative 2D gel Southern analysis of joint molecules from *ECM11-AID* subcultures with and without the addition of auxin while maintaining pachytene arrest. **d,** Quantification of total joint molecule levels from experiment represented in panel **c**. **e, f,** Representative images of surface-spread meiotic nuclei from pachytene-arrested *ECM11-AID* cells sampled 1 hr after the addition of auxin or DMSO vehicle at 7 hrs, and immunostained for the indicated markers. Scale bars = 1 μm. **g,** Quantification of Zip3 and Msh5 immunostaining foci from the *ECM11-AID* experiments represented in panels **e** and **f**, and comparison with corresponding data from *ZIP1-AID* (from **Fig. 4f**). 40-60 nuclei were counted in each case. Error bars represent SD. Unpaired two-tailed *t* test, \*\*\*\**P*<0.0001. **h, i,** Quantification of colocalization between immunostaining foci of Zip1 and Zip3 (**h**), and Zip1 and Msh5 (**i**) from auxin-treated *ECM11-AID* nuclei represented in panels **e** and **f** with auxin treated cells. Error bars represent SD.

Immunostaining revealed that SCs disassembled following Ecm11-AID degradation in arrested *NDT80-IN* cells, although numerous foci of Zip1 persisted (**Fig. 6e**). Thus, the central element is required to both establish and maintain full synapsis. Msh5 and Zip3 foci were maintained at ∼50% of no-auxin control levels (**Fig. 6f,g**), in contrast to the almost complete loss seen following degradation of Rec8-AID, Zip1-AID, or Msh4-AID. Moreover, remaining Msh5 and Zip3 foci showed high degrees of colocalization with residual Zip1 structures (**Fig. 6h,i**). These observations suggest that while mature SC may stabilize a subset of CRCs, local ensembles of Zip1 and CRCs can be sufficient to protect dHJs and support crossover-specific resolution.

## Discussion

The SC is an ancient structure that evolved with the emergence of sexual reproduction. Several functions are attributed to the SC: assembly of homolog axes and pairwise synapsis of homologs establish topological order in the nucleus; synapsis also attenuates formation of DSBs and facilitates their repair, thereby diminishing DNA-damage checkpoint signaling; and at designated crossover sites, SC components facilitate dHJ formation^5^. This study reveals that key SC components, but not necessarily end-to-end synapsis *per se*, also enable crossover-specific resolution of dHJs by protecting them from unscheduled dissolution into noncrossovers by the conserved BLM/STR complex (**Extended Data Fig. 1b,c**). Failure of this critical final step of meiotic recombination will result in unlinked univalent chromosomes that are prone to missegregation, resulting in aneuploid gametes that are associated with infertility, miscarriage, and congenital disease in humans.

Our data indicate that dHJs formed at crossover-designated recombination sites do not possess an intrinsic structure or topology that constrains their resolution fate to ensure crossing over. We previously proposed a model of crossover-specific resolution in which asymmetric loading of PCNA during the DNA synthesis associated with dHJ formation subsequently directs strand-specific nicking by the MutLγ endonuclease on both sides of the two Holliday junctions, but distal to the exchange points^4^. This pattern of incisions always specifies a crossover outcome when the nicked dHJs are disassembled by the BLM/STR complex. This two-step resolution model reconciles the counterintuitive pro-crossover role of BLM/STR during meiosis, which contradicts its well-characterized anti-crossover roles in unwinding D-loops and dissolving unnicked dHJs into noncrossver products (**Extended Data Figs. 1b and 3b,c**).

Within this framework, we propose that mature dHJs are unnicked and therefore vulnerable to STR-catalyzed dissolution until MutLγ is activated via Ndt80-dependent expression of Plk1^Cdc5^. By preventing BLM/STR from acting on dHJs until they have been nicked by MutLγ, SC components impose the sequential steps required for crossover-specific resolution (**Extended Data Fig. 1b**).

We suggest that local ensembles of cohesin and SC transverse filaments can be sufficient protect dHJs by stabilizing CRCs, components of which can directly bind HJs (including MutSγ, MutLγ, Zip2^SHOC1^-Spo16, and Mer3^HFM1^)^24,40^ and may directly compete with BLM/STR for binding dHJs and/or may constrain its activity. This suggestion is consistent with electron-microscopy visualization of late recombination nodules associated with short patches of SC in *Sordaria macrospora*^41^; and with the inference that, in *C. elegans,* the cohesin local to designated crossover sites is distinctly regulated^27^. Moreover, components of the SC-central region help recruit and stabilize CRCs ^31,42–45^. Also in *C. elegans*, SC-central region proteins assemble transient crossover-specific compartments or “bubbles” that may protect CRCs until crossover-specific resolution is executed^43^.

We also found that CRCs are required to maintain synapsis during pachytene. This observation is consistent with studies indicating that the SC central region is initially dynamic and labile but becomes more stable, contingent upon the development of CRCs^46–49^. Intriguingly, new subunits are continually incorporated into the SC central region after synapsis^46,49^; and in budding yeast, incorporation of new Zip1 molecules occurs predominantly at CRC sites^46^. Given that Zip1 appears to recruit CRC components directly^44,50^, this suggests a possible mechanism for the mutual stabilization of SCs and CRCs. This relationship may result in SCs with non-uniform structure and stability^46^, which could influence resolution fates and explain why sites of crossing over are the last to desynpase during diplotene^51^. The interdependence and dynamicity of SCs and CRCs will render chromosomal interactions readily reversible until dHJs are resolved into crossovers. These attributes could help adjust and proof-read homologous synapsis, minimize and resolve synaptic interlocks^52^, and preserve genome stability by destabilizing interactions between non-allelic and diverged sequences.

Importantly, dependency between synapsis and CRCs likely helps impose the ordered sequence of events required to ensure crossing over between homologs. At designated crossover sites, CRCs promote and maintain homolog synapsis, facilitate the formation of dHJs, and protect them from aberrant dissolution. In turn, synapsis globally attenuates DSB formation^53^, diminishing the DNA-damage kinase signaling that inhibits Ndt80 activity. Ndt80-dependent expression of Plk1^Cdc5^ then activates crossover-specific dHJ resolution, CRCs disassemble, and desynapsis ensues (**Fig. 1c,d**)

Our analysis also reveals that two SMC complexes mediate essentially all Plk1^Cdc5^-dependent joint-molecule resolution during meiosis (**Extended Data Fig. 1c**). Rec8-cohesin is required to maintain synapsis and CRCs, and thereby protects dHJs to facilitate crossover-specific dHJ resolution. Whether local or global functions of Rec8-cohesin are important for crossover-specific dHJ resolution, and the roles of cohesive versus chromatin-loop organizing populations of Rec8-cohesin remain unclear. An independent Smc5/6 pathway is essential for resolution by Mus81-Mms4^EME1^ and a subpopulation of BLM/STR, producing a mixture of crossovers and noncrossovers. Despite these distinctions, Rec8-cohesin and Smc5/6 could have common functions to constrain favorable joint-molecule topology^54^ and control access by resolving enzymes.

## Methods

### Data reporting

No statistical methods were used to predetermine sample size. The experiments were not randomized, and the investigators were not blinded to allocation during experiments. Blinding was employed during outcome assessment for cytology experiments.

### Yeast Strains

For full genotypes see **Supplementary Information Table 1**. The AID system ^10^ was optimized for meiosis by replacing the promoter of the *P_ADH1_-OsTIR1* cassette with the *CUP1* promoter^35^. C-terminal fusion of a minimal AID degron to targeted proteins was constructed using plasmid pHyg-AID*-9Myc as template for PCR-mediated allele replacement ^55^. To construct an internal degron allele of *ZIP1,* AID degron sequences were inserted into plasmid pMPY-3xHA and integrated after codon 700 via PCR epitope tagging ^56^. Primers used to construct AID degron alleles are listed in **Supplementary Information Table 2**. The estrogen-inducible *IN-NDT80 GAL4-ER* system has been described ^57–59^.

### Meiotic Time Courses and DNA Physical Assays

Detailed protocols for meiotic time courses and DNA physical assays at the *HIS4::LEU2* locus have been described ^60^. At 6.5 h after induction of meiosis, CuSO4 (100 mM stock in dH2O) was added for a final concentration of 50 μM to induce expression of *P_CUP1_-OsTIR1* (encoding the Tir1 E3 ligase) and cell cultures were split. At 7 h, estradiol (5 mM stock, Sigma E2758 in ethanol) was added for a final concentration of 1 μM to both subcultures to induce *NDT80-IN*. Simultaneously, auxin (3-indoleacetic acid, Sigma 13750, 2 M stock in DMSO) was added to one subculture for a final concentration of 2 mM; an equivalent volume of DMSO was added to the no-auxin control subculture. At 7.5 h, auxin was added again at 1 mM. To analyze the timing and efficiency of meiotic divisions and sporulation, cells were fixed in 40% ethanol, 0.1 M sorbitol, stained with DAPI, and ∼200 cells were categorized for each time point. For imaging, DAPI-stained cells were mounted in antifade (Vectashield, Vector Laboratories, Inc.) and digital images captured using a Zeiss AxioPlan II microscope, Hamamatsu ORCA-ER CCD camera and Volocity software.

### Chromosome spreading and immunofluorescence microscopy

Cell samples were collected and processed for chromosome spreading and immunostaining essentially as described ^61^. Primary antibodies kindly provided by Akira Shinohara were chicken anti-Red1 (1:500 dilution), rabbit anti-Msh5 (1:750), and rabbit anti-Zip3 (1:500); anti-Zip1 (1:400) was a gift from Scott Keeney; anti-myc (1:1000, Roche 11667149001) was used to detect AID-9myc fused proteins. All primary antibodies were incubated overnight at room temperature in 100 μl TBS/BSA buffer (10 mM Tris pH 7.5, 150 mM NaCl, 1% BSA).

Secondary antibodies anti-rabbit 568 (A11036 Molecular Probes, 1:1,000), anti-mouse 488 (A11029 Molecular Probes, 1:1,000), anti-rabbit 647 (A21245 Invitrogen), and anti-guinea pig 555 (A21435 Life Technologies) were incubated for 1 h at 37 °C. Coverslips were mounted with Prolong Gold antifade reagent (Invitrogen, P36930). Digital images were captured using a Zeiss Axioplan II microscope, Hamamatsu ORCA-ER CCD camera and analyzed using Volocity software. Scatterplots were generated using the GraphPad program in Prism.

### Western Blot Analysis

Whole cell extracts were prepared using a TCA extraction method, essentially as described ^62^. Samples were analyzed by standard SDS-PAGE and Western blotting using the following primary antibodies: anti-c-Myc (1:1000, Roche 11667149001), anti-HA (1:1000, Sigma11583816001), Arp7 (1:10,000, Santa Cruz SC-8960), anti-Msh4 (1:500), and anti-Msh5 (1:500, Msh4 and Msh5 antibodies were a gift from Dr. Akira Shinohara). Secondary antibodies (1:5000) were IRDye® 800CW Donkey anti-Mouse IgG (LI-COR 925-32212), IRDye® 680LT Donkey anti-Goat IgG (LI-COR 925-68024), IRDye® 680LT Donkey anti-Rabbit IgG (LI-COR 925-68023) and IRDye® 800CW Donkey anti-Rabbit IgG (LI-COR 925-32213). Western blots were imaged on an Odyssey Infrared Imager (LI-COR) and quantification of protein bands was performed using Image Studio Lite Ver 4.0 software.

### Statistical analysis and Reproducibility

Statistical analyses were performed using Prism (GraphPad software Inc.). For bar graphs and scatter plots comparing two samples (aggregated from three or more replicate experiments), unpaired *t*-tests were performed. For multiple comparisons, one-way ANOVA was performed (Tukey or Dunnett tests, depending on the specific comparisons being made). For scatter plots and bar graphs, error bars show the mean value with standard deviation. Western blots are representative of at least three repeats (Figs. 1b, 4a 4b, Fig. 5c and 5d; and Extended Data Figs. 1a, 2a, 3a, 3c, 3e, 4a, 4g, 6a, 6f, 7a, 7g). Representative immunofluorescence images were chosen from over 60 samples analyzed in each of at least three biological replicates (Figs. 1e, h, 2a, 3c, 2e, 2g, 3a, 4e 4g, 5a, 5e, 6a, 6c and 6e).

## Data availability

Relevant data generated or analyzed during this study are included in this Article and its Supplementary Information files. Biological materials are available from the corresponding author.

## End Notes

### Acknowledgements

We thank A. Shinohara and S. Keeney for reagents; J. Matos for communicating unpublished data; and members of the Hunter laboratory for support and discussions. S.H. was supported by the predoctoral Training Program in Molecular and Cellular Biology at UC Davis supported by NIH T32 training grant GM007377. This work was supported by NIH NIGMS grant GM074223 to N.H. N.H. is an Investigator of the Howard Hughes Medical Institute.

## Author contributions

S.M.T. and N.H. conceived the study and designed the experiments. S.H., R.B., J.E.M., J.K., M.P., E.N., N.L., C.M., H.L., and M.L. assisted with experiments and data analysis. S.M.T and N.H. wrote the manuscript with inputs and edits from all authors.

## Additional Information

**Supplementary Information** is available for this paper.

**Correspondence and requests for materials** should be addressed to N.H.

**Reprints and permissions information** is available at www.nature.com/reprints.

**Extended Data Fig. 1.**
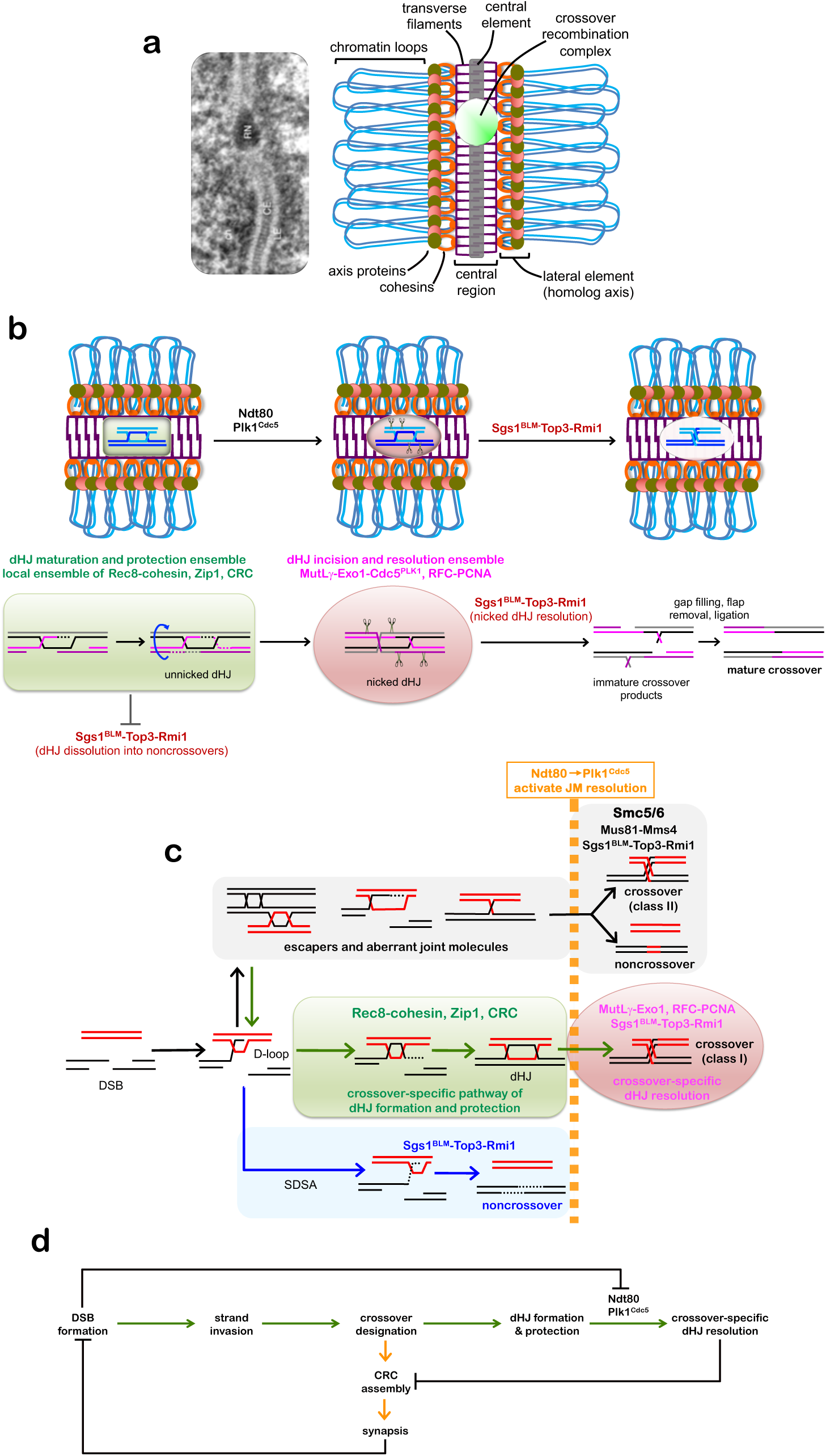
Summary and Model. **a**, Electron micrograph of SC from *Blaps cribrosa*^63^ and schematic highlighting the key features of pachytene-stage chromosomes. Ch, chromatin; LE, lateral element; CE, central element, RN, recombination nodule (the site of a crossover recombination complex, CRC). **b**, Rec8-cohesin, Zip1, and CRCs constitute local dHJ maturation and protection ensembles that facilitates dHJ formation and protects unnicked dHJs from being dissolved into non-crossovers by the BLM/STR complex. dHJ protection must be maintained until Ndt80-dependent expression of polo-like kinase Plk1^Cdc^^5^ triggers crossover-specific resolution via a two-step mechanism: (i) strand-specific nicking of dHJs by the MutLγ endonuclease and associated factors^4^; (ii) resolution of nicked dHJs by the BLM/STR complex. Placement of the nicks, on both sides of the two Holliday junctions but distal to the exchange points, specifies a crossover outcome. **c**, Pathways of joint-molecule resolution during meiotic recombination. Resected DSB ends undergo homology search and DNA strand exchange to form nascent displacement loops (D-loops) as common precursors to all pathways. Non-crossover fated D-loops are unwound by BLM/STR to effect synthesis-dependent strand annealing (SDSA), independently of Plk1^C^^dc^^5^. D-loops that are designated a crossover fate mature into dHJs. In this class-I pathway, Plk1^Cdc^^5^ triggers two-step crossover-specific resolution as described in **b**. All other joint-molecule structures are resolved via a Smc5/6-dependent pathway that is also dependent on Plk1^Cdc^^5^ and yields class-II crossovers and non-crossovers. **d**, Dependencies that help ensure crossing over between homologs. At designated crossover sites, CRC assembly promotes and maintains homolog synapsis and dHJs. In turn, synapsis globally attenuates DSB formation^53^, thereby diminishing the DNA-damage kinase signaling that inhibits Ndt80 activity^64^. Ndt80-dependent expression of Plk1^Cdc^^5^ then activates crossover-specific dHJ resolution, CRCs disassemble, and desynapsis ensues.

**Extended Data Fig. 2.**
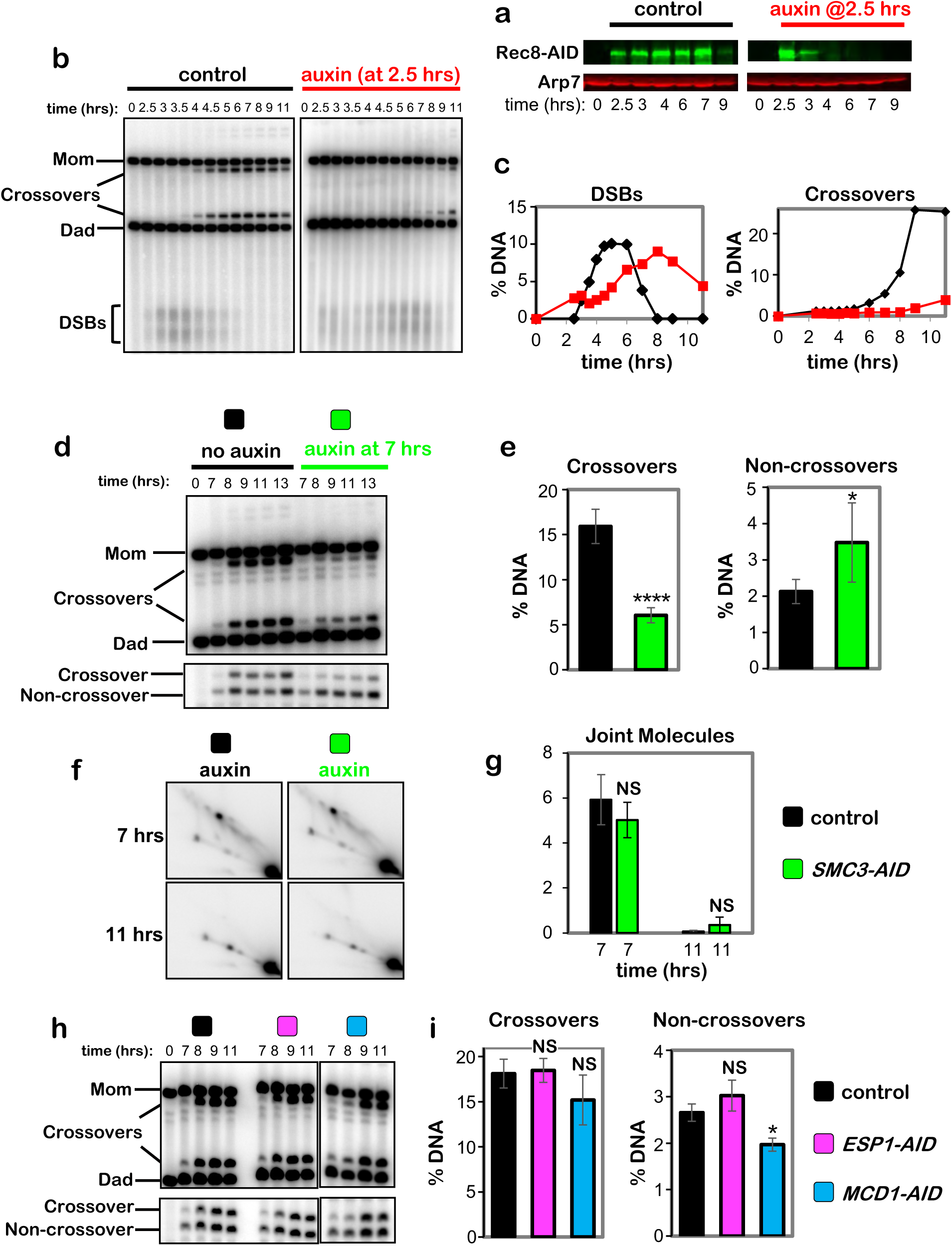
Early functions of Rec8 and roles of Smc3, Kleisin Mdc1^Rad^^21^, and Separase Esp1 in crossover-specific dHJ resolution. **a**, Western analysis of early Rec8-AID degradation following addition of auxin at 2.5 hrs. **b**, 1D-gel Southern analysis of DSBs and crossovers following early degradation of Rec8-AID. **c**, Quantification of the DSB and crossover level from the Southern blot shown in C. **d**, Representative 1D-gel Southern analysis of crossover (upper panel) and non-crossover (lower panel) formation from *SMC3-AID* subcultures with or without auxin. **e**, Quantification of final levels of crossovers and non-crossovers at 13 hrs from *SMC3-AID* subcultures with or without auxin (mean ± SD, 4 independent experiments, \*\*\*\**P*<0.0001, NS, not significant *P*=0.0561, two-tailed unpaired *t*-test). **f**, Representative 2D gel Southern analysis of joint molecules from *SMC3-AID* subcultures with or without auxin, before and after release from arrest at 7 and 11 hrs, respectively. **g**, Quantification of total joint molecule levels from experiments represented in panel f (mean ± SD, 3 independent experiments, NS, not significant, *P*=0.3167 for levels at 7hrs; NS, not significant, *P* >0.9999 for levels at 11hrs; two-tailed unpaired *t*-test). **h**, Representative 1D-gel Southern analysis of crossover (upper panel) and non-crossover (lower panel) formation in control, *ESP1-AID* (with auxin) or *MCD1-AID* (with auxin) strains. **i**, Final levels of crossovers and non-crossovers at 11 hrs from the indicated strains (mean ± SD, 3 independent experiments, \**P*=0.0146; NS, not significant, *P*=0.9610 (crossovers, control vs. *ESP1-AID*), *P*=0.1793 (crossovers, control vs. *MCD1-AID*), *P*=0.2073 (noncrossovers, control vs. *ESP1-AID*), Dunnett’s multiple comparisons test, control).

**Extended Data Fig. 3.**
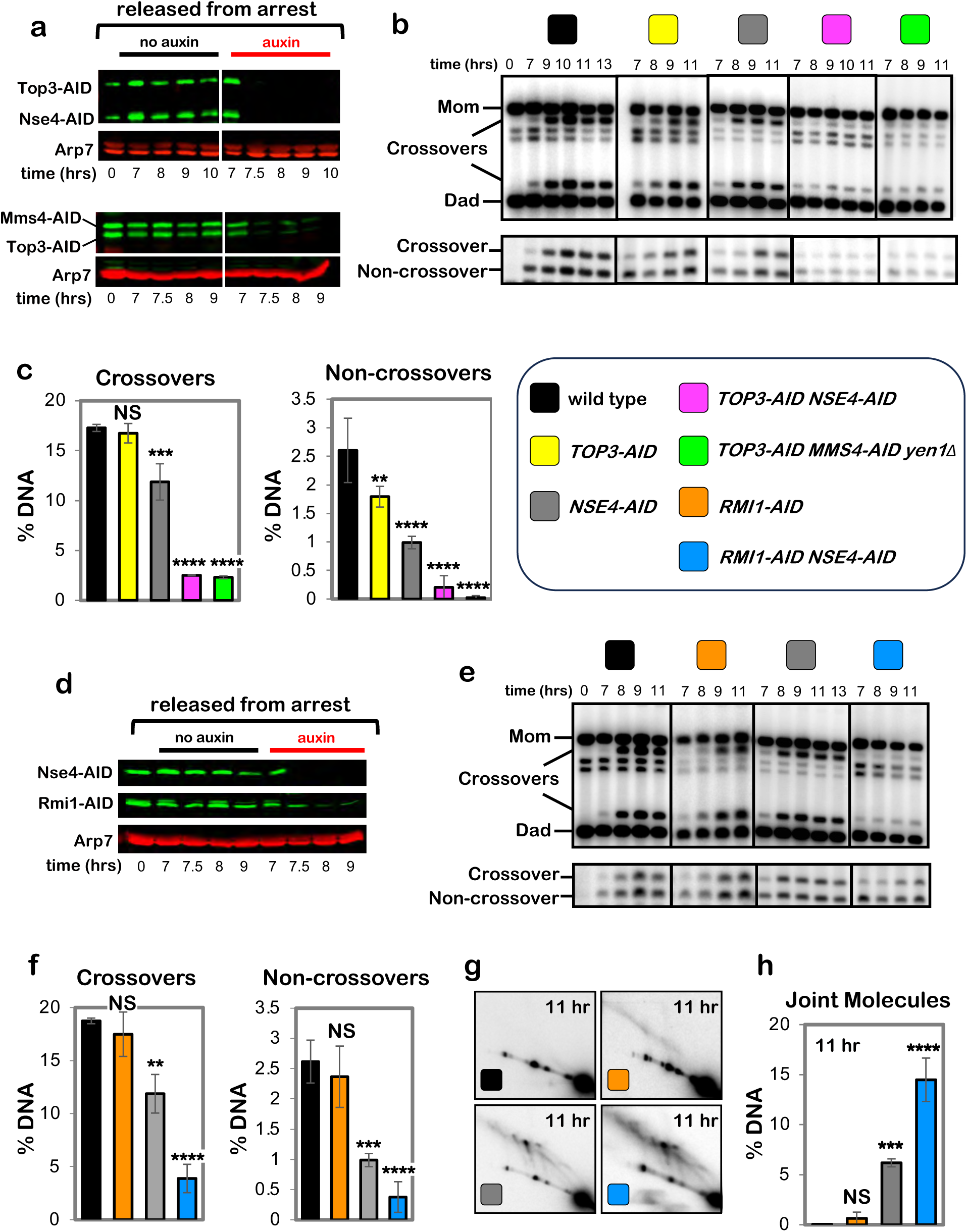
Sgs1-Top3-Rmi1 and Smc5/6 mediate essentially all joint-molecule resolution. **a**, Western blot analysis of Top3-AID and Nse4-AID co-degradation in cells released from pachytene-arrest and addition of auxin at 7 hours. **b**, Representative 1D-gel Southern analysis of crossover (upper panel) and non-crossover (lower panel) formation in control, *TOP3-AID* (with auxin), *NSE4-AID* (with auxin), *TOP3-AID NSE4-AID* (with auxin) and *TOP3-AID NSE4-AID yen1Δ* (with auxin) following release from pachytene arrest. **c**, Final levels of crossovers and non-crossovers at 11 hrs from experiments represented in **b** (mean ± SD). Statistical comparisons with control: \*\*\*\**P*<0.0001, \*\*\**P*=0.0003, NS, not significant, *P*=0.3101 (crossovers, *TOP3-AID* vs. control), *P*=0.003 (crossovers, *NSE4-AID* vs. control), *P*= 0.0029 (non-crossovers *TOP3-AID* vs. control), Dunnett’s multiple comparisons test. Number of experiments: control: control: n=3, *TOP3-AID*: n=3, *NSE4-AID*: n=3, *TOP3-AID NSE4-AID*: n=3 and *TOP3-AID NSE4-AID yen1Δ*, n=2. **d**, Western blot analysis of RMI1-AID and Nse4-AID co-degradation in cells released from pachytene-arrest and addition of auxin at 7 hours. **e**, Representative 1D-gel Southern analysis of crossover (upper panel) and non-crossover (lower panel) formation in control, *RMI1-AID* (with auxin), *NSE4-AID* (with auxin) and *RMI1-AID NSE4-AID* (with auxin) following release from pachytene arrest. **f**, Final levels of crossovers and non-crossovers at 11 hrs from experiments represented in **b** (mean ± SD). Statistical comparisons with control: \*\*\*\**P*<0.0001, \*\*\**P*=0.0002, \*\**P*=0.0016, NS, not significant, *P*=0.9928 (crossovers, *RMI1*-*AID* vs. control), *P*=0.7036 (non-crossovers *RMI1-AID* vs. control), Dunnett’s multiple comparisons test. Number of experiments: control: control: n=4, *RMI1-AID*: n=2, *NSE4-AID*: n=4 and *RMI1-AID NSE4-AID*: n=3. **g**, Representative 2D gel Southern analysis of joint molecules from control, *RMI1-AID, NSE4-AID and RMI1-AID NSE4-AID* subcultures with auxin after release from arrest at 11 hrs. **h**, Quantification of total joint molecule levels from experiments represented in panel **g** (mean ± SD), NS, not significant, *P*=0.8596, \*\*\**P*=0.0002, \*\*\*\**P*<0.0001, Dunnett’s multiple comparisons test. control: n=3, *RMI1-AID*: n=3, *NSE4-AID*: n=3 and *RMI1-AID NSE4-AID*: n=3.

**Extended Data Fig. 4.**
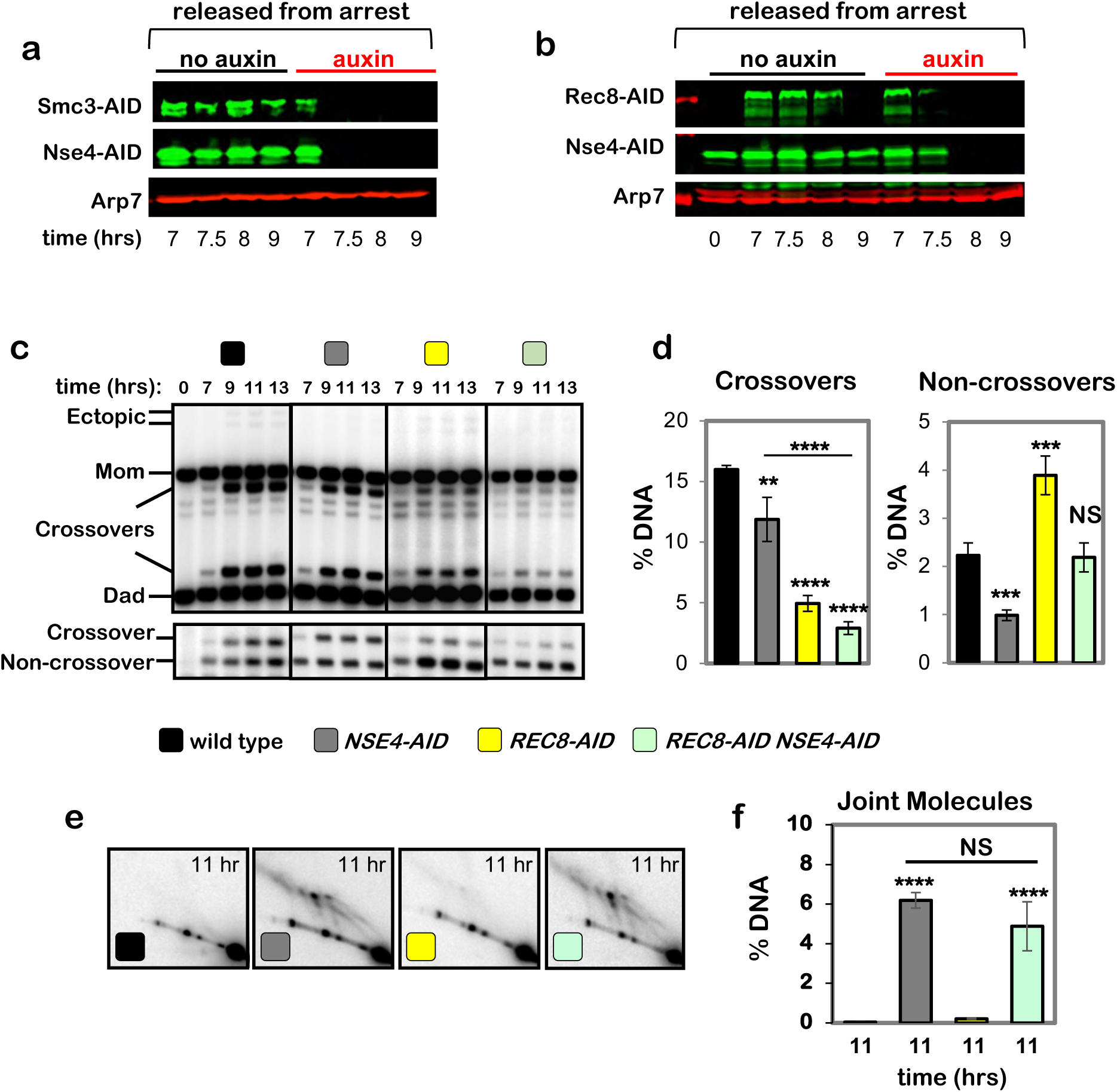
Cohesin and the Smc5/6 complex act in parallel pathways of joint-molecule resolution. **a**, Western blot images of Smc3-AID Nse4-AID co-degradation following auxin addition at 7 hours (corresponds to experiments in Fig2. e–g). Arp7 is a loading control. **b**, Western blot images of Rec8-AID Nse4-AID co-degradation following auxin addition at 7 hours. **c**, Representative 1D-gel Southern analysis of crossover (upper panel) and non-crossover (lower panel) formation in control, *NSE4-AID* (with auxin), *REC8-AID* (with auxin), and *NSE4-AID REC8-AID* (with auxin) strains. **d**, Final levels of crossovers and non-crossovers at 11 hrs from experiments represented in **c** (mean ± SD, 3 independent experiments). Statistical comparisons with control unless indicated. Dunnett’s multiple comparisons test, \*\*\*\**P*<0.0001, \*\**P*=0.0049 (*NSE4-AID* vs. control), \*\*\**P*=0.0002 (*REC8-AID* vs. control), \*\*\**P*=0.0022 (*NSE4-AID* vs. control), NS, not significant *P*=0.9955). **e**, Representative Southern blot 2D gel images at 11 hours from no auxin control, *NSE4-AID* (with auxin), *REC8-AID* (with auxin) and *REC8-AID NSE4-AID* (with auxin). f, Quantification of total joint-molecule levels from the experiments represented in e (mean ± SD, 3 independent experiments). NS, not significant, *P*=0.3320 Tukey’s multiple comparison test.

**Extended Data Fig. 5.**
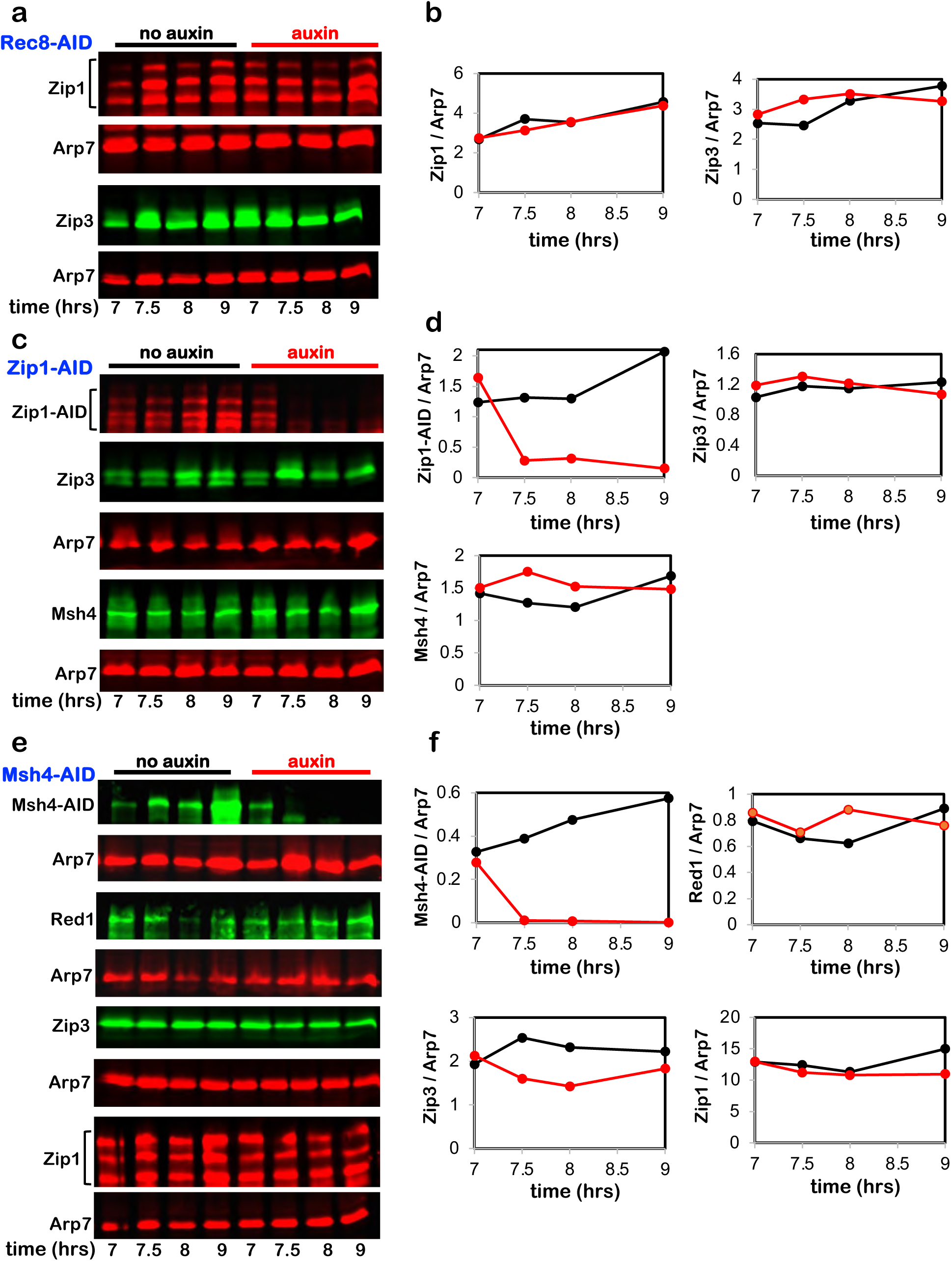
Targeted degradation does not cause off-target effects. **a**, Representative Western blot analysis of Zip1 and Zip3 with or without Rec8-AID degradation following auxin addition at 7 hours. **b**, Quantification of Zip1 and Zip3 level from the experiments shown in a. **c**, Representative Western blot analysis of Zip3 and Msh4 with or without Zip1-AID degradation following auxin addition at 7 hours. **d**, Quantification of Zip1-AID, Zip3, and Msh4 levels from the experiments shown in c. **e**, Representative Western blot analysis of Red1, Zip3, and Zip1 with or without Msh4-AID degradation following auxin addition at 7 hours. **F**, Quantification of Msh4-AID, Red1, Zip3, and Zip1 level from the experiments shown in e. Arp7 is the loading control in all experiments.

**Extended Data Fig. 6.**
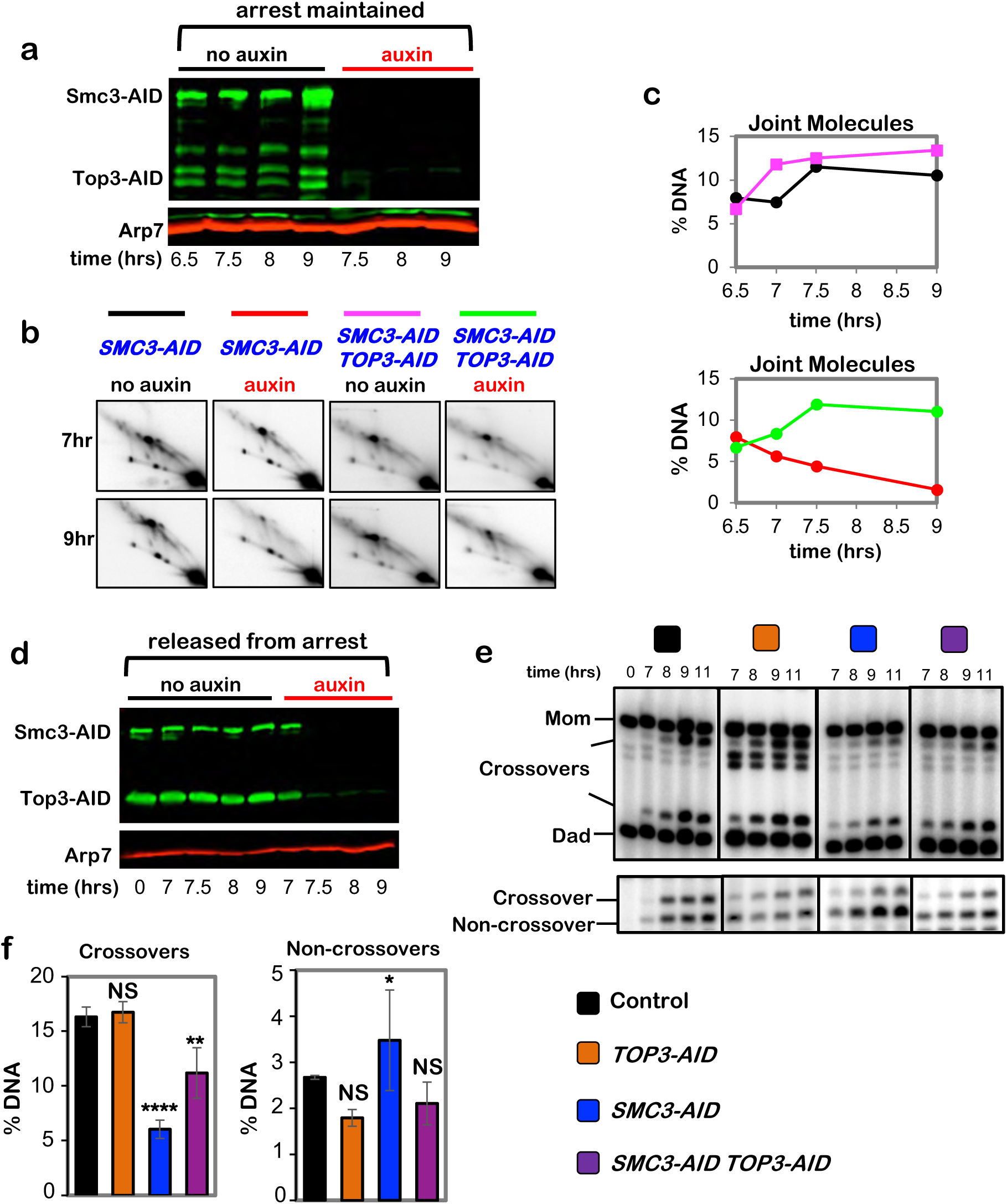
Cohesin protects dHJs from dissolution by the BLM/STR complex. **a**, Western blot analysis of Smc3-AID and Top3-AID co-degradation in pachytene-arrested cells following the addition of auxin at 7 hours. Arp7 is a loading control. **b**, Representative 2D gel Southern analysis of joint molecules from subcultures of pachytene arrested *SMC3-AID* and *SMC3-AID TOP3-AID* strains, with or without the addition of auxin. **c**, Quantification of total joint molecules from experiments shown in b. **d**, Western blot analysis of Smc3-AID and Top3-AID co-degradation in cells released from pachytene-arrest and addition of auxin at 7 hours. Arp7 is used as loading control. **e**, Representative 1D-gel Southern analysis of crossover (upper panel) and non-crossover (lower panel) formation in control, *TOP3-AID* (with auxin), *SMC3-AID* (with auxin) and *SMC3-AID TOP3-AID* (with auxin) following release from pachytene arrest. **f**, Final levels of crossovers and non-crossovers at 11 hrs from experiments represented in **g** (mean ± SD, 3 (*SMC3-AID* and *TOP3-AID SMC3-AID*) or 4 (*TOP3-AID* and *SMC3-AID*) independent experiments). Statistical comparisons with control: \*\*\*\**P*<0.0001, ** *P*= 0.0038, NS, not significant, *P*=0.9876 (crossovers, *TOP3-AID* vs. control), \**P*=0.0423. NS, not significant, *P*=0.8549 (non-crossovers, *TOP3-AID* vs. control), NS, not significant, *P* >0.9999 (non-crossovers, *TOP3-AID SMC3-AID* vs. control), Dunnett’s multiple comparisons test.

**Extended Data Fig. 7.**
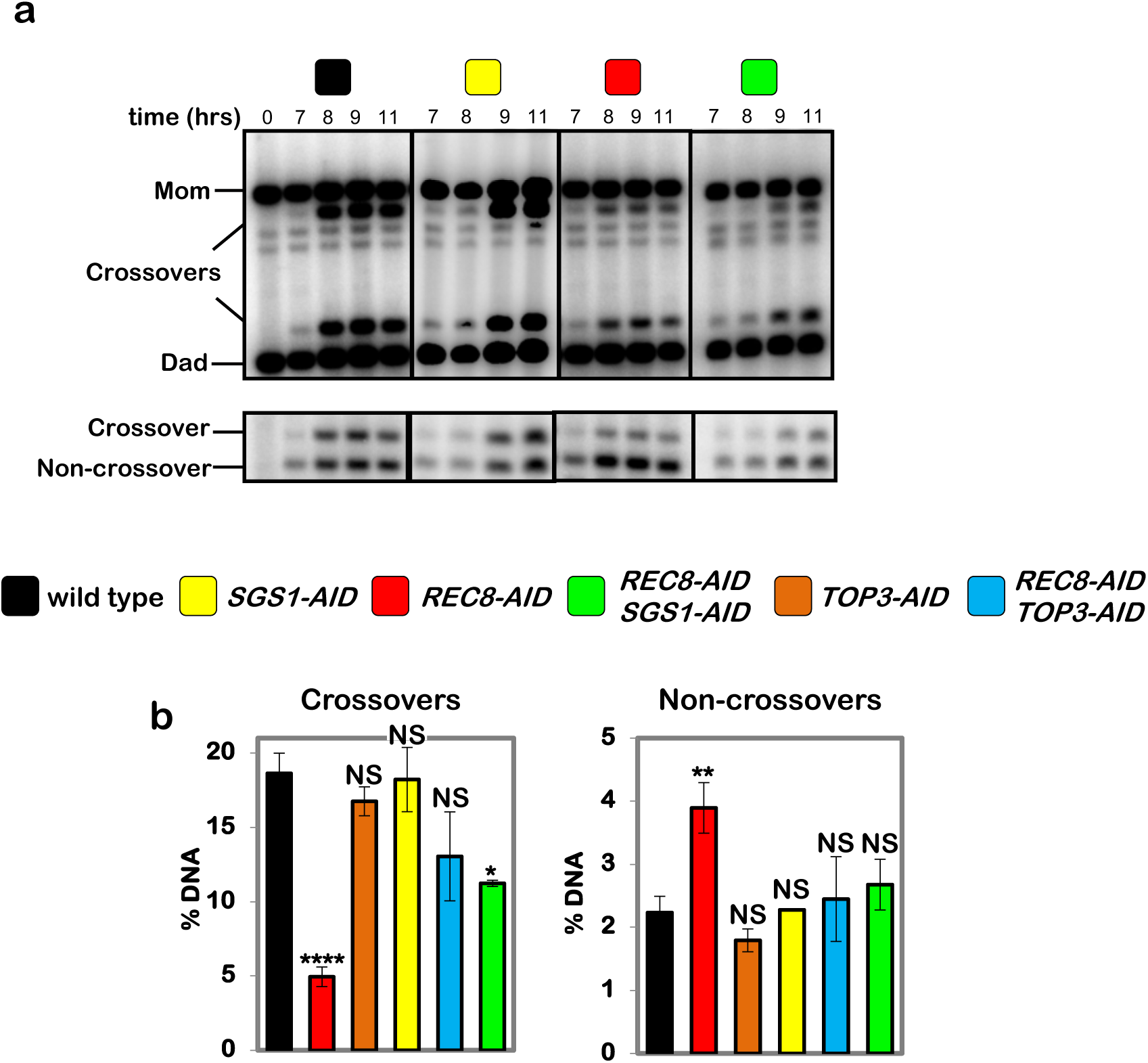
Sgs1-AID degradation partially suppresses the crossover defect resulting from degradation of Rec8-AID. **a**, Representative 1D-gel Southern analysis of crossover (upper panel) and non-crossover (lower panel) formation following release from pachytene arrest in control, *SGS1-AID* (with auxin), *REC8-AID* (with auxin), *REC8-AID SGS1-AID* (with auxin). **b**, Final levels of crossovers and non-crossovers at 11 hrs from the indicated strains (mean ± SD). Data for *TOP3-AID* and *REC8-AID TOP3-AID* from **Fig. 5e** are shown for comparison. \*\*\*\**P*<0.0001, \**P*=0.0162, \*\**P*=0.0012; NS, not significant. Crossovers: *P*=0.9378 (*TOP3-AID* vs. control), *P*=0.3345 (*SGS1-AID* vs. control), *P*=0.1490 (*REC8-AID TOP3-AID* vs. control); Non-crossovers: *P*=0.4759 (*TOP3-AID* vs. control), *P*>0.9999 (*SGS1-AID* vs. control), *P*= 0.9465 (*REC8-AID TOP3-AID* vs. control), *P*=0.5646 (*REC8-AID SGS1-AID* vs. control), Dunnett’s multiple comparisons tests. Number of experiments: control n=3, *TOP3-AID* n=3, *SGS1-AID* n=2, *REC8-AID* n=3, REC8-AID SGS1-AID n=2, *TOP3-AID REC8-AID* n=2.

**Extended Data Fig. 8.**
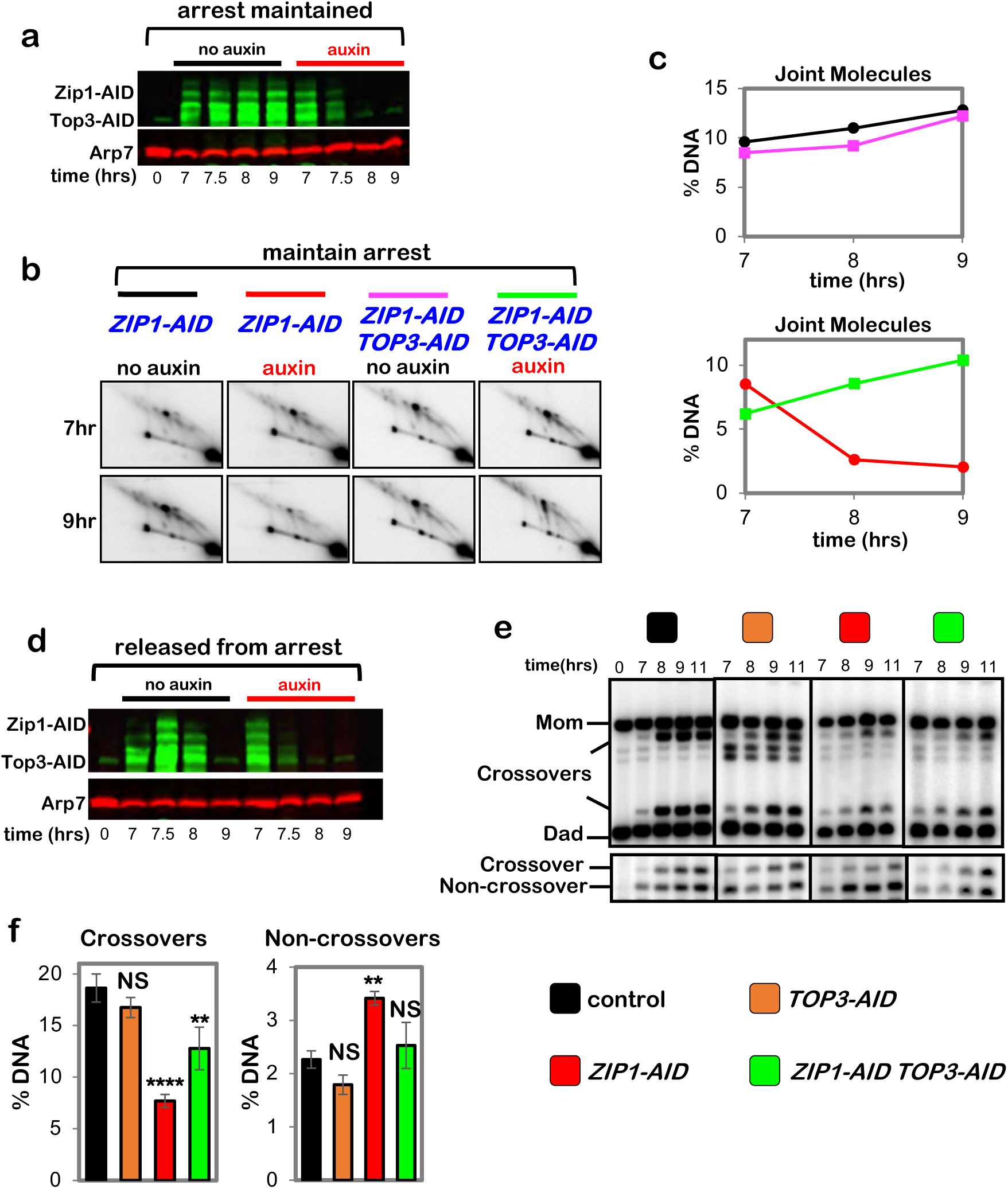
Zip1 protects double-Holliday junctions from aberrant resolution mediated by the BLM/STR complex. **a**, Western blot analysis of Zip1-AID and Top3-AID co-degradation in pachytene-arrested cells following the addition of auxin at 7 hours. Arp7 is a loading control. **b**, Representative 2D gel Southern analysis of joint molecules from subcultures of pachytene arrested *ZIP1-AID* and *ZIP1-AID TOP3-AID* strains, with or without the addition of auxin. **c**, Quantification of total joint molecules from experiments shown in b. **d**, Western blot analysis of Zip1-AID and Top3-AID co-degradation in cells released from pachytene-arrest and addition of auxin at 7 hours. **e**, Representative 1D-gel Southern analysis of crossover (upper panel) and non-crossover (lower panel) formation in control, *TOP3-AID* (with auxin), *ZIP1-AID* (with auxin) and *ZIP1-AID TOP3-AID* (with auxin) following release from pachytene arrest. **f**, Final levels of crossovers and non-crossovers at 11 hrs from experiments represented in **g** (mean ± SD). Statistical comparisons with control: \*\*\*\**P*<0.0001, \*\**P*=0.0022, NS, not significant. *P*=0.1011 (crossovers *TOP3-AID* vs. control), *P*=0.1104 (non-crossovers *TOP3-AID* vs. control), *P*=0.5171 (non-crossovers *ZIP1-AID TOP3-AID* vs. control), Dunnett’s multiple comparisons tests. Number of experiments: control: n=3, *TOP3-AID*: n=3, *ZIP1-AID*: n=2, *ZIP1-AID TOP3-AID*: n=2.

**Extended Data Fig. 9.**
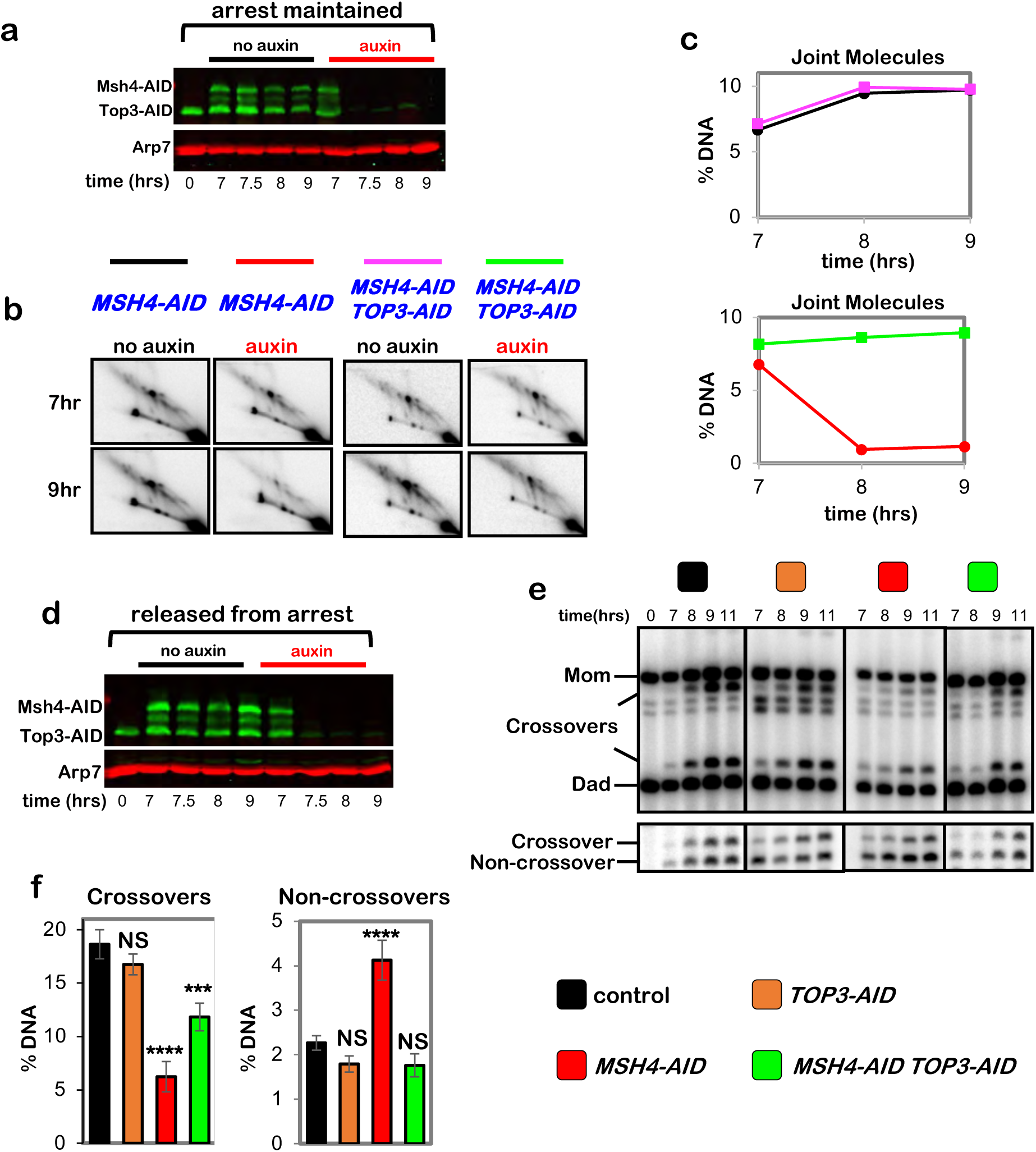
MutSγ protects double-Holliday junctions from aberrant resolution mediated by the BLM/STR complex. **a**, Western blot analysis of Msh4-AID and Top3-AID co-degradation in pachytene-arrested cells following the addition of auxin at 7 hours. Arp7 is a loading control. **b**, Representative 2D gel Southern analysis of joint molecules from subcultures of pachytene arrested *MSH4-AID* and *MSH4-AID TOP3-AID* strains, with or without the addition of auxin. **c**, Quantification of total joint molecules from experiments shown in b. **d**, Representative 1D gel Southern analysis of non-crossover formation from subcultures of pachytene arrested *MSH4-AID* and *MSH4-AID TOP3-AID* strains, with or without the addition of auxin (the same experiments analyzed in panel b). **e**, Quantification of non-crossovers from pachytene arrested *MSH4-AID* (red) and *MSH4-AID TOP3-AID* (green) strains following the addition of auxin. **f**, Western blot analysis of Msh4-AID and Top3-AID co-degradation in cells released from pachytene-arrest and addition of auxin at 7 hours. **g**, Representative 1D-gel Southern analysis of crossover (upper panel) and non-crossover (lower panel) formation in control, *TOP3-AID* (with auxin), *MSH4-AID* (with auxin) and *MSH4-AID TOP3-AID* (with auxin) following release from pachytene arrest. **h**, Final levels of crossovers and non-crossovers at 11 hrs from experiments represented in **g** (mean ± SD). Statistical comparisons with control: \*\*\*\**P*<0.0001, \*\*\**P*=0.0003, NS, not significant, *P*>0.9999 (crossovers, *TOP3-AID* vs. control), *P*=0.0821 (non-crossovers *TOP3-AID* vs. control), *P*=0.0638 (non-crossovers *TOP3-AID MSH4-AID* vs. control), Dunnett’s multiple comparisons test. Number of experiments: control: control: n=6, *TOP3-AID*: n=3, *MSH4-AID*: n=6, *TOP3-AID MSH4-AID*: n=3.

**Extended Data Fig. 10.**
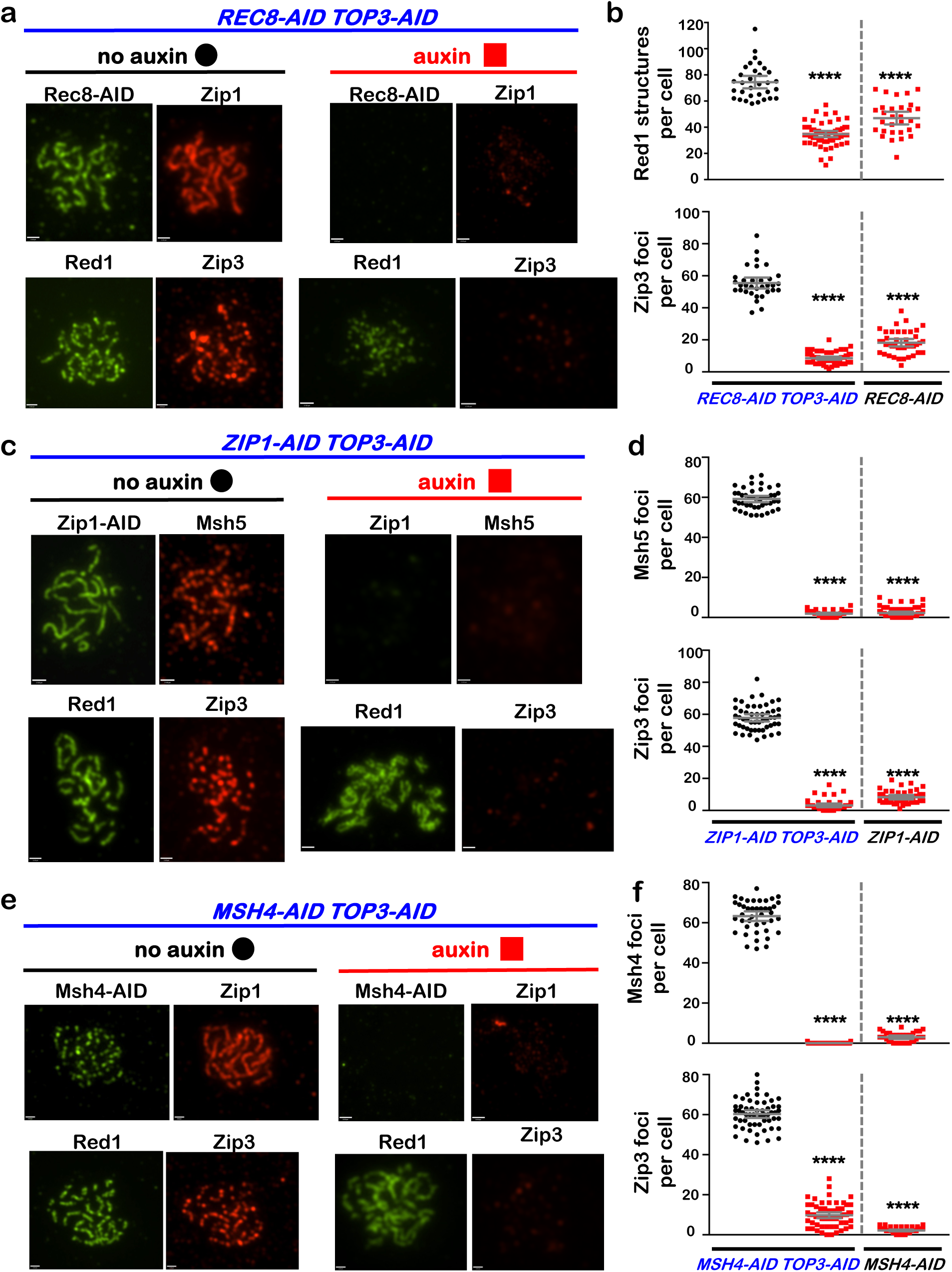
Stabilizing dHJs in pachytene does not bypass interdependence between cohesin, synapsis, and crossover recombination complexes. **a**, Representative images of surface-spread meiotic nuclei from pachytene-arrested *REC8-AID TOP3-AID* cells sampled 1 hr after the addition of auxin or DMSO vehicle at 7 hrs, and immunostained for the indicated markers. **b**, Quantification of Red1 and Zip3 immunostaining structures from the *REC8-AID TOP3-AID* experiments represented in panel a, and comparison with corresponding data from *REC8-AID* (from Fig. 3b). **c**, Representative images of surface-spread meiotic nuclei from pachytene-arrested *ZIP1-AID TOP3-AID* cells sampled 1 hr after the addition of auxin or DMSO vehicle at 7 hrs, and immunostained for the indicated markers. **d**, Quantification of Msh5 and Zip3 immunostaining foci from the experiments represented in panel c, and comparison with corresponding data from *ZIP1-AID* (from Fig. 4f). **e**, Representative images of surface-spread meiotic nuclei from pachytene-arrested *MSH4-AID TOP3-AID* cells sampled 1 hr after the addition of auxin or DMSO vehicle at 7 hrs, and immunostained for the indicated markers. **f**, Quantification of Msh4 and Zip3 immunostaining foci from the experiments represented in panel e, and comparison with corresponding data from *MSH4-AID* (from Fig. 4h). Scale bars = 1 μm. Error bars represent SD. 40-60 nuclei were counted in each case. Unpaired two-tailed *t* test, \*\*\*\**P*<0.0001.

