## Supplemental Tables 1-2 for "Protection of double-Holliday junctions ensures crossing over during meiosis"

| Supplementary Table 1. *Saccharomyces cerevisiae* strains used in this study | |
|---|---|
| Strain | Genotype |
| NHY 7291 | *MATa/MATα HIS4::LEU2-(BamHI)/his4-X::LEU2-(NgoMIV)—URA3 TOP3-AID-9myc::hphMX4 NSE4-AID-9myc::hphMX4 /TOP3-AID-9myc::hphMX4 NSE4-AID-9myc::hphMX4 hphMX4::PGAL1-NDT80/hphMX4::PGAL1-NDT80 ura3:PGPD1-GAL4(848)-ER:URA3 hphMX4::PCUP1-1-OsTIR1::lys2/ PCUP1-1-OsTIR1-9Myc-URA3* |
| NHY7475 | *MATa/MATα HIS4::LEU2-(BamHI)/his4-X::LEU2-(NgoMIV)—URA3 NSE4-AID-9myc::hphMX4 /NSE4-AID-9myc::hphMX4 hphMX4::PGAL1-NDT80/hphMX4::PGAL1-NDT80 ura3:PGPD1-GAL4(848)-ER:URA3 hphMX4::PCUP1-1-OsTIR1::lys2/ PCUP1-1-OsTIR1-9Myc-URA3* |
| NHY7699 | *MATa/MATα HIS4::LEU2-(BamHI)/his4-X::LEU2-(NgoMIV)—URA3 REC8-AID-9myc::hphMX4 NSE4-AID-9myc::hphMX4 /REC8-AID-9myc::hphMX4 NSE4-AID-9myc::hphMX4 hphMX4::PGAL1-NDT80/hphMX4::PGAL1-NDT80 ura3:PGPD1-GAL4(848)-ER:URA3 hphMX4::PCUP1-1-OsTIR1::lys2/ PCUP1-1-OsTIR1-9Myc-URA3* |
| NHY7824 | *MATa/MATα HIS4::LEU2-(BamHI)/his4-X::LEU2-(NgoMIV)—URA3 REC8-AID-9myc::hphMX4 /REC8-AID-9myc::hphMX4 hphMX4::PGAL1-NDT80/hphMX4::PGAL1-NDT80 ura3:PGPD1-GAL4(848)-ER:URA3 hphMX4::PCUP1-1-OsTIR1::lys2/ PCUP1-1-OsTIR1-9Myc-URA3* |
| NHY7854 | *MATa/MATα HIS4::LEU2-(BamHI)/his4-X::LEU2-(NgoMIV)—URA3 NSE4-AID-9myc::hphMX4 csm2Δ::KanMX6 /NSE4-AID-9myc::hphMX4 csm2Δ::KanMX6 hphMX4::PGAL1-NDT80/hphMX4::PGAL1-NDT80 ura3:PGPD1-GAL4(848)-ER:URA3 hphMX4::PCUP1-1-OsTIR1::lys2/ PCUP1-1-OsTIR1-9Myc-URA3* |
| NHY7914 | *MATa/MATα HIS4::LEU2-(BamHI)/his4-X::LEU2-(NgoMIV)—URA3 NSE4-AID-9myc::hphMX4 mph1::KanMX6/NSE4-AID-9myc::hphMX4 mph1::KanMX6 hphMX4::PGAL1-NDT80/hphMX4::PGAL1-NDT80 ura3:PGPD1-GAL4(848)-ER:URA3 hphMX4::PCUP1-1-OsTIR1::lys2/ PCUP1-1-OsTIR1-9Myc-URA3* |
| NHY7988 | *MATa/MATα HIS4::LEU2-(BamHI)/his4-X::LEU2-(NgoMIV)—URA3 REC8-AID-9myc::hphMX4 mlh3Δ::kanMX4 /REC8-AID-9myc::hphMX4 mlhΔ::kanMX4 hphMX4::PGAL1-NDT80/hphMX4::PGAL1-NDT80 ura3:PGPD1-GAL4(848)-ER:URA3 hphMX4::PCUP1-1-OsTIR1::lys2/ PCUP1-1-OsTIR1-9Myc-URA3* |
| NHY8111 | *MATa/MATα HIS4::LEU2-(BamHI)/his4-X::LEU2-(NgoMIV)—URA3 REC8-AID-9myc::hphMX4 TOP3-AID-9myc::hphMX4/REC8-AID-9myc::hphMX4 TOP3-AID-9myc::hphMX4 hphMX4::PGAL1-NDT80/hphMX4::PGAL1-NDT80 ura3:PGPD1-GAL4(848)-ER:URA3 hphMX4::PCUP1-1-OsTIR1::lys2/ PCUP1-1-OsTIR1-9Myc-URA3* |
| NHY8177 | *MATa/MATα HIS4::LEU2-(BamHI)/his4-X::LEU2-(NgoMIV)—URA3 TOP3-AID-9myc::hphMX4 /TOP3-AID-9myc::hphMX4 hphMX4::PGAL1-NDT80/hphMX4::PGAL1-NDT80 ura3:PGPD1-GAL4(848)-ER:URA3 hphMX4::PCUP1-1-OsTIR1::lys2/ PCUP1-1-OsTIR1-9Myc-URA3* |
| NHY8256 | *MATa/MATα HIS4::LEU2-(BamHI)/his4-X::LEU2-(NgoMIV)—URA3 SMC3-AID-9myc::hphMX4 /SMC3-AID-9myc::hphMX4 hphMX4::PGAL1-NDT80/hphMX4::PGAL1-NDT80 ura3:PGPD1-GAL4(848)-ER:URA3 hphMX4::PCUP1-1-OsTIR1::lys2/ PCUP1-1-OsTIR1-9Myc-URA3* |
| NHY8263 | *MATa/MATα HIS4::LEU2-(BamHI)/his4-X::LEU2-(NgoMIV)—URA3 SMC3-AID-9myc::hphMX4 NSE4-AID-9myc::hphMX4 /SMC3-AID-9myc::hphMX4 NSE4-AID-9myc::hphMX4 hphMX4::PGAL1-NDT80/hphMX4::PGAL1-NDT80 ura3:PGPD1-GAL4(848)-ER:URA3 hphMX4::PCUP1-1-OsTIR1::lys2/ PCUP1-1-OsTIR1-9Myc-URA3* |
| NHY8500 | *MATa/MATα HIS4::LEU2-(BamHI)/his4-X::LEU2-(NgoMIV)—URA3 SMC3-AID-9myc::hphMX4 TOP3-AID-9myc::hphMX4 /SMC3-AID-9myc::hphMX4 TOP3-AID-9myc::hphMX4 hphMX4::PGAL1-NDT80/hphMX4::PGAL1-NDT80 ura3:PGPD1-GAL4(848)-ER:URA3 hphMX4::PCUP1-1-OsTIR1::lys2/ PCUP1-1-OsTIR1-9Myc-URA3* |
| NHY8555 | *MATa/MATα HIS4::LEU2-(BamHI)/his4-X::LEU2-(NgoMIV)—URA3 MMS4-AID-9myc::hphMX4 /MMS4-AID-9myc::hphMX4 hphMX4::PGAL1-NDT80/hphMX4::PGAL1-NDT80 ura3:PGPD1-GAL4(848)-ER:URA3 hphMX4::PCUP1-1-OsTIR1::lys2/ PCUP1-1-OsTIR1-9Myc-URA3* |
| NHY8714 | *MATa/MATα HIS4::LEU2-(BamHI)/his4-X::LEU2-(NgoMIV)—URA3 SMC3-AID-9myc::hphMX4 MMS4-AID-9myc::hphMX4 yen1Δ::KanMX6 / SMC3-AID-9myc::hphMX4 MMS4-AID-* |

| | |
|---|---|
| | *9myc::hphMX4 yen1Δ::KanMX6 hphMX4::PGAL1-NDT80/hphMX4::PGAL1-NDT80 ura3:PGPD1-GAL4(848)-ER:URA3 hphMX4::PCUP1-1-OsTIR1::lys2/ PCUP1-1-OsTIR1-9Myc-URA3* |
| NHY8793 | *MATa/MATα HIS4::LEU2-(BamHI)/his4-X::LEU2-(NgoMIV)—URA3 SGS1-AID-9myc::hphMX4 NSE4-AID-9myc::hphMX4 /SGS1-AID-9myc::hphMX4 NSE4-AID-9myc::hphMX4 hphMX4::PGAL1-NDT80/hphMX4::PGAL1-NDT80 ura3:PGPD1-GAL4(848)-ER:URA3 hphMX4::PCUP1-1-OsTIR1::lys2/ PCUP1-1-OsTIR1-9Myc-URA3* |
| NHY8806 | *MATa/MATα HIS4::LEU2-(BamHI)/his4-X::LEU2-(NgoMIV)—URA3 SGS1-AID-9myc::hphMX4 /SGS1-AID-9myc::hphMX4 hphMX4::PGAL1-NDT80/hphMX4::PGAL1-NDT80 ura3:PGPD1-GAL4(848)-ER:URA3 hphMX4::PCUP1-1-OsTIR1::lys2/ PCUP1-1-OsTIR1-9Myc-URA3* |
| NHY8873 | *MATa/MATα HIS4::LEU2-(BamHI)/his4-X::LEU2-(NgoMIV)—URA3 TOP3-AID-9myc::hphMX4 MMS4-AID-9myc::hphMX4 yen1Δ::KanMX6 /TOP3-AID-9myc::hphMX4 MMS4-AID-9myc::hphMX4 yen1Δ::KanMX6 hphMX4::PGAL1-NDT80/hphMX4::PGAL1-NDT80 ura3:PGPD1-GAL4(848)-ER:URA3 hphMX4::PCUP1-1-OsTIR1::lys2/ PCUP1-1-OsTIR1-9Myc-URA3* |
| NHY8875 | *MATa/MATα HIS4::LEU2-(BamHI)/his4-X::LEU2-(NgoMIV)—URA3 MMS4-AID-9myc::hphMX4 yen1Δ::KanMX6 /MMS4-AID-9myc::hphMX4 yen1Δ::KanMX6 hphMX4::PGAL1-NDT80/hphMX4::PGAL1-NDT80 ura3:PGPD1-GAL4(848)-ER:URA3 hphMX4::PCUP1-1-OsTIR1::lys2/ PCUP1-1-OsTIR1-9Myc-URA3* |
| NHY8880 | *MATa/MATα HIS4::LEU2-(BamHI)/his4-X::LEU2-(NgoMIV)—URA3 TOP3-AID-9myc::hphMX4 NSE4-AID-9myc::hphMX4 mph1::KanMX6 /TOP3-AID-9myc::hphMX4 NSE4-AID-9myc::hphMX4 mph1::KanMX6 hphMX4::PGAL1-NDT80/hphMX4::PGAL1-NDT80 ura3:PGPD1-GAL4(848)-ER:URA3 hphMX4::PCUP1-1-OsTIR1::lys2/ PCUP1-1-OsTIR1-9Myc-URA3* |
| NHY9078 | *MATa/MATα HIS4::LEU2-(BamHI)/his4-X::LEU2-(NgoMIV)—URA3 ZIP1-AID-3HA /ZIP1-AID-3HA hphMX4::PGAL1-NDT80/hphMX4::PGAL1-NDT80 ura3:PGPD1-GAL4(848)-ER:URA3 hphMX4::PCUP1-1-OsTIR1::lys2/ PCUP1-1-OsTIR1-9Myc-URA3* |
| NHY9117 | *MATa/MATα HIS4::LEU2-(BamHI)/his4-X::LEU2-(NgoMIV)—URA3 RMI1-AID-9myc::hphMX4 NSE4-AID-9myc::hphMX4 /RMI1-AID-9myc::hphMX4 NSE4-AID-9myc::hphMX4 hphMX4::PGAL1-NDT80/hphMX4::PGAL1-NDT80 ura3:PGPD1-GAL4(848)-ER:URA3 hphMX4::PCUP1-1-OsTIR1::lys2/ PCUP1-1-OsTIR1-9Myc-URA3* |
| NHY9263 | *MATa/MATα HIS4::LEU2-(BamHI)/his4-X::LEU2-(NgoMIV)—URA3 ECM11-AID-9myc::hphMX4 /ECM11-AID-9myc::hphMX4 hphMX4::PGAL1-NDT80/hphMX4::PGAL1-NDT80 ura3:PGPD1-GAL4(848)-ER:URA3 hphMX4::PCUP1-1-OsTIR1::lys2/ PCUP1-1-OsTIR1-9Myc-URA3* |
| NHY9347 | *MATa/MATα HIS4::LEU2-(BamHI)/his4-X::LEU2-(NgoMIV)—URA3 MSH4-AID-9myc::hphMX4 /MSH4-AID-9myc::hphMX4 hphMX4::PGAL1-NDT80/hphMX4::PGAL1-NDT80 ura3:PGPD1-GAL4(848)-ER:URA3 hphMX4::PCUP1-1-OsTIR1::lys2/ PCUP1-1-OsTIR1-9Myc-URA3* |
| NHY9525 | *MATa/MATα HIS4::LEU2-(BamHI)/his4-X::LEU2-(NgoMIV)—URA3 TOP3-AID-9myc::hphMX4 ZIP1-AID-3HA /TOP3-AID-9myc::hphMX4 ZIP1-AID-3HA hphMX4::PGAL1-NDT80/hphMX4::PGAL1-NDT80 ura3:PGPD1-GAL4(848)-ER:URA3 hphMX4::PCUP1-1-OsTIR1::lys2/ PCUP1-1-OsTIR1-9Myc-URA3* |
| NHY9554 | *MATa/MATα HIS4::LEU2-(BamHI)/his4-X::LEU2-(NgoMIV)—URA3 MSH4-AID-9myc::hphMX4 TOP3-AID-9myc::hphMX4/MSH4-AID-9myc::hphMX4 TOP3-AID-9myc::hphMX4 hphMX4::PGAL1-NDT80/hphMX4::PGAL1-NDT80 ura3:PGPD1-GAL4(848)-ER:URA3 hphMX4::PCUP1-1-OsTIR1::lys2/ PCUP1-1-OsTIR1-9Myc-URA3* |
| * In addition, all strains contain the markers *leu2::hisG, ura3(Δsma-pst)* and *ho::hisG* | |

### Supplementary Table 2. Oligonucleotides used in this study

| | |
|---|---|
| REC8-AID primer 1 | TGCGGTAACTAAAGATCTTAAATTGAGAAGAGAGGACGAAATAATTGTATATGCCCGTACGCTGCAGGTCGAC |
| REC8-AID primer 2 | GAGGACAGCGGCTAGTAACCGCTGTCCTCATATGGAAGGAGAAAATAAAAATTCAATCGATGAATTCGAGCTCG |
| NSE4-AID primer1 | GGACCAGCTATGAAAAAAAAAAAAAAAAAAAAAAAAAAAACTGTACATATTATATGCAGCGCTCTATCGCTGTTAATCGATGAATTCGAGCTCG |
| NSE4-AID primer1 | ATTATTTTTCAAATGGACATGCCTACTTGGCGAAAACTAATAAAGAAATACAACATCACTTCACCATTCTTAGACCGTACGCTGCAGGTCGAC |
| TOP3-AID primer1 | GAATGCCTGCAAGAATACTCTCTTGCAAGTTTATGACCGTGTCAAGGCGTCCATGCGTACGCTGCAGGTCGAC |
| TOP3-AID primer2 | TCATGCAATTAAGCGGAGGGCTTTTTCGAAGACAAAAGGCGGCAAAAACGCCTTAATCGATGAATTCGAGCTCG |
| MMS4-AID primer1 | GAGGCAGTAGAAAAAAGATTGTACAAACTGTTTACTTGTACTGATCCAAATGATACTATTGAACGTACGCTGCAGGTCGAC |
| MMS4-AID Primer2 | GCAGTGATTTTCAAAACGACTGCCTTAAGGTATGTTCTTATATACAAAGTTTCGTTCGATCATCAATCGATGAATTCGAGCTCG |
| SGS1-AID Primer1 | TGCTAATGGGAGACGAGGTTTTAGAAATTACCGAGGTCACTATAGAGGAAGAAAGCGTACGCTGCAGGTCGAC |
| SGS1-AID Primer2 | GCTTGGCGAATGGTGTCGTAGTTATAAGTAACACTATTTATTTTTCTACTCTTCAATCGATGAATTCGAGCTCG |
| ZIP1 internal AID Primer1 | GAGGAATCACTAAGCGATGTAAAAACCCTAAAACAGCAAGTGATAGTTTTGAAATCGGAGAAGCAAGATATAACAAGGGAACAAAAGCTGGAG |
| ZIP1 interal primer2 | GTAAATTTTTGGTGACTTCTTCCAAACTTTCGAGGTTATCTTGAAGTTCTAACTTTTCGGCGCCACCTCCGCCTCCACCCTGTAGGGCGAATTGGGTAC |
| SMC3-AID Primer1 | GTTATTGAGGTCAATAGAGAAGAAGCAATCGGATTCATTAGAGGTAGCAATAAATTCGCTGAAGTCCGTACGCTGCAGGTCGAC |
| SMC3-AID Primer2 | CAAATAGCTATTTATGTAAGCAAAACTGATATTTTTATATACAAACCGTTTCAAATATCTCTTAATCGATGAATTCGAGCTCG |
| MCD1-AID Primer1 | GTCAAACAGAAGCATTCGGAAATATTAAAATAGACGCCAAACCTGCACTATTTGAAAGGTTTATCAATGCTCGTACGCTGCAGGTCGAC |
| MCD1-AID Primer2 | GTCTTTGATCTATATATGCATCAGCTTACTGGGTCCACCAAGAAATCCCCTCGGCGTAACTAGGTTTTAATCGATGAATTCGAGCTCG |
| ESP1-AID Primer1 | GAAGTGTATGCCATCTACGTTACTTGAACGGCGCAGCTCCTGTTATTTATGGGTTACCGATCAAGTTCGTATCACGTACGCTGCAGGTCGAC |
| ESP1-AID Primer2 | CAAAATCGGATTTCCCATGCTTTTTCTCAATGTCTATATGAAATCTTTTCGAAACAACCAGTACATGTAACAATTAATCGATGAATTCGAGCTCG |
| MSH4-AID Primer1 | GGAAATGAAAAAGAGCCCTTGACTTTAGGGAAATTAAAAGAAATAAACTCCGACTTCATCGAAAATTTTGAAGAACGTACGCTGCAGGTCGAC |
| MSH4-AID Primer2 | CATTTTCTCCGGTTTTTGATAACTCTGTACAGAAATAATGGATTATAGTTTTAAGCTAAGCGGAAAAGCCAAATTAATCGATGAATTCGAGCTCG |
